# A Bayesian framework reveals heterogeneous and stochastic decision-making in NK cell cytotoxicity

**DOI:** 10.64898/2026.09.16.752048

**Authors:** Elephes Sung, Cathal Hosty, Khodor S. Hazime, Leanne Peiser, Lara Stepan, Daniel M. Davis, Ruben Perez-Carrasco

## Abstract

Natural killer (NK) cells display striking variability in their cytotoxic responses, but whether this variation reflects random events, stable differences between cells, or effects of previous encounters remains unclear. We developed a Bayesian framework that uses single-cell interaction histories to disentangle these sources of variability. The framework models target encounters and killing decisions to quantify population heterogeneity, determine the sample sizes needed to distinguish competing mechanisms, and separate stable cell-to-cell differences from history-dependent behaviour. Synthetic data established when these mechanisms can be reliably distinguished in practice. We tested the framework using timelapse imaging of NK cells exposed to rituximab or CC-96673, an antibody co-targeting CD20 and CD47. Despite similar overall killing, the framework identified distinct underlying responses: rituximab mainly increased mean killing rate, whereas CC-96673 reduced cell-to-cell variation. Event-count analysis supported continuous population heterogeneity, while ordered interaction histories revealed that both stable differences in killing propensity and previous encounters shape cytotoxic decisions. To make the framework directly usable, we introduce BARRACUDA, an open-source web platform and Python package implementing these analyses, including donor-aware extensions, for single-cell cytotoxicity datasets.

## 1 Introduction

At the single cell level, natural killer (NK) cell cytotoxicity is a multiscale process emerging from the interplay of cell migration, target engagement, immunological synapse formation, delivery of lethal hits, and subsequent detachment [1]. Considerable progress has been made in characterising many of these components individually. For example, extensive work has dissected the molecular processes in immunological synapse assembly, structure, duration and function [2–4]. Similarly, single cell omics approaches have revealed extensive phenotypic diversity across cytotoxic lymphocyte populations [5, 6]. These approaches, however, primarily provide static or population level snapshots and therefore do not directly capture the dynamic sequence of interactions through which individual NK cells engage, kill, and move between target cells over time.

To address this, time-lapse imaging has become a powerful tool for studying NK cell cytotoxicity both *in vitro* and *in vivo* [7–10]. By tracking individual cells, these experiments provide temporal profiles of cell migration, interaction, and cytotoxic decision-making. One striking observation is the prevalent heterogeneity in cytotoxic decision-making and contact-kill histories among individual NK cells. This observation has motivated attempts to classify NK cells into behavioural subpopulations. For instance, Vanherberghen et al. classified primary NK cells into killing, non-killing, exhausted, and stochastic subsets based on their interaction histories [9]. However, such classifications remain largely descriptive. Even when behavioural classes can be predicted from imaging data using machine learning approaches [11], the mechanistic origins of cytotoxic variability remain unresolved.

A central challenge is to identify the mechanisms that give rise to the heterogeneity observed in NK cell contact-kill trajectories. Variability in cytotoxic behaviour can emerge from multiple sources, including stochastic target encounters, stochastic killing decisions, stable cell-to-cell differences in cytotoxic capacity, systematic variation between donors, or behavioural changes induced by previous interactions. Disentangling these contributions from time-lapse imaging data remains difficult, and systematic frameworks for separating them are limited [12, 13]. This challenge is compounded by the fact that distinct biological mechanisms can generate similar contact-kill trajectories. As a result, it remains unclear how much of the observed variability reflects intrinsic cellular heterogeneity, how much emerges from interaction history, and how much is simply a consequence of stochasticity. This ambiguity becomes particularly apparent in serial-killing NK cells, the subset of the population that repeatedly contact and eliminate multiple targets during longitudinal observation [9, 14–17]. The emergence of these highly cytotoxic cells can be explained by fundamentally different mechanisms. They may simply represent the tail of a stochastic process, a “lucky” subset of otherwise identical NK cells that experienced particularly favourable sequences of encounters and killing events. Alternatively, they may correspond to intrinsically distinct cells with enhanced cytotoxic capacity, compatible with the molecular variability observed in their gene expression profile [6]. A third possibility is that cytotoxic behaviour evolves during the interaction process itself, such that previous encounters modify the probability of future killing events. Consistent with this idea, NK cells can switch from fast granzyme-B-mediated killing during early encounters to slower death-receptor-mediated killing during later encounters [18], while sequential stimulation through different NK cell receptors can produce order-dependent responses [19]. In practice, all three mechanisms are likely to operate simultaneously. The central question is therefore how much information about their relative contributions is encoded within contact-kill trajectories and whether these contributions can be quantitatively disentangled.

Here, we develop a quantitative mathematical framework to infer the mechanistic origins of cytotoxic variability from single cell interaction histories. We first focus on disentangling variability arising from stochastic target encounters and stochastic killing decisions, allowing the heterogeneity profiles of NK cell populations to be quantified and the experimental sample sizes required to distinguish competing models of cellular heterogeneity to be determined. We then extend the framework to analyse ordered contact-kill trajectories, enabling us to test whether cytotoxic variability is an intrinsic property of individual NK cells or an emergent consequence of their interaction history. Throughout the study, we combine synthetic datasets to characterise inference performance under different mechanistic scenarios with time-lapse imaging data from NK cells exposed to rituximab or CC-96673. CC-96673 (BMS-986358) is a bispecific antibody co-targeting CD20 and CD47. Its high affinity CD20 binding arm directs the antibody to CD20^+^ target cells, and its detuned affinity CD47 binding arm blocks the inhibitory CD47–SIRP*α* interaction [20]. Despite their different mechanisms of action, rituximab and CC-96673 produced similar overall levels of killing. Therefore, it provides an opportunity to investigate whether similar population-level outcomes can emerge from distinct underlying single-cell mechanisms.

To make the framework broadly accessible, we also introduce BARRACUDA, an open-source implementation of the framework developed in this study. The web application supports interactive analysis of event counts and ordered contact histories, with donor-aware models available for count data. The accompanying cyto-barracuda Python package supports reproducible scripted analyses and integration into existing workflows. Together, these interfaces make the methods developed here available for application to further single-cell cytotoxicity datasets.

## 2 Results

### 2.1 A Bayesian framework for inferring the origins of cytotoxic variability

The challenge addressed in this study is fundamentally an inverse problem: how much mechanistic information is encoded within contact-kill trajectories? Bayesian inference provides a natural framework for addressing this problem by evaluating experimental observations against the probabilistic mathematical predictions of different hypotheses (likelihood). The result of this inference process is a posterior (credibility) distribution, containing the plausibility of alternative mechanisms and their associated parameters. This allows us to ask not only which biological explanations are consistent with the data, but also whether the available observations are sufficient to distinguish between competing hypotheses.

A key advantage of this probabilistic formulation is that different biological mechanisms can be incorporated as additional layers within the likelihood function. Stochasticity, cellular heterogeneity, and history-dependent behaviour can each be represented as separate model components, allowing hypotheses to be nested and combined within a common inference framework. This enables progressively more complex mechanistic models to be constructed and quantitatively compared against experimental observations, providing a systematic route for dissecting the origins of cytotoxic variability.

Each NK history extracted from time-lapse microscopy can be encoded as a time-ordered sequence of binary contact outcomes. For example, the trajectory *x*_*i*_ = (0, 1, 0, 0) represents a cell that performed four contacts in total: an initial non-lethal contact, followed by a successful killing event, and two subsequent non-lethal contacts. Different levels of information can be extracted from these trajectories. At the coarsest level, one can consider only the total number of contacts performed by each cell, *N*_*i*_, corresponding to the total number of interaction events recorded in the trajectory. A second summary statistic is the total number of successful killing events 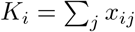 which ignores the ordering of events but retains information about cytotoxic output.

### 2.2 Event-count models quantify stochasticity versus heterogeneity in the population

The framework introduced below is general and can be applied to any discrete event recorded in a timelapse trajectory. For clarity, we first consider the number of contact events per trajectory, where *N*_*i*_ denotes the number of target encounters performed by NK cell *i* during the observation period. The same analysis will later be applied to the population distribution of killing events *K*_*i*_.

We begin with the simplest hypothesis in which the variability observed across NK cell populations can be explained by stochasticity alone. Under this hypothesis, all NK cells are assumed to behave identically, and differences in the number of observed contacts or kills arise solely from the random timing of individual events. In other words, all cells have the same probability per unit time of engaging with or killing a target cell. Some cells may appear highly cytotoxic simply because they experience a favourable sequence of stochastic encounters and killing decisions, rather than because they possess intrinsically different cytotoxic capabilities.

For bulk chamber inter-cellular interactions, we assume that the contact rate between NK cells and tumour cells remains constant longitudinally within the observation window. Under this assumption, the number of contacts accumulated by each NK cell follows a Poisson process. In this scenario, the distribution of observed contact counts is fully specified by a single parameter, the event rate *λ*, representing the expected number of contacts during the observation period. We label this model as ℳ_homo_,

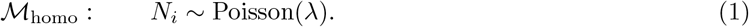

Deviations from the Poisson prediction have long been interpreted as evidence of underlying population heterogeneity [21]. Here, rather than simply detecting such deviations, we seek to quantify both the magnitude and structure of this heterogeneity. To do so, we extend the stochastic model into a hierarchical framework in which each individual NK cell possesses its own event rate *λ*_*i*_, drawn from a population-level distribution. This distribution captures a continuum of behaviours, ranging from rarely interacting cells to highly active cells. We further allow for the possibility of a subpopulation of non-engaging cells whose event rate remains effectively zero throughout the observation period. Together, these assumptions define a zero-inflated Gamma model for the cell-specific rates (see Fig. 1) that we label as ℳ_ZIΓ_,

**Fig. 1.**
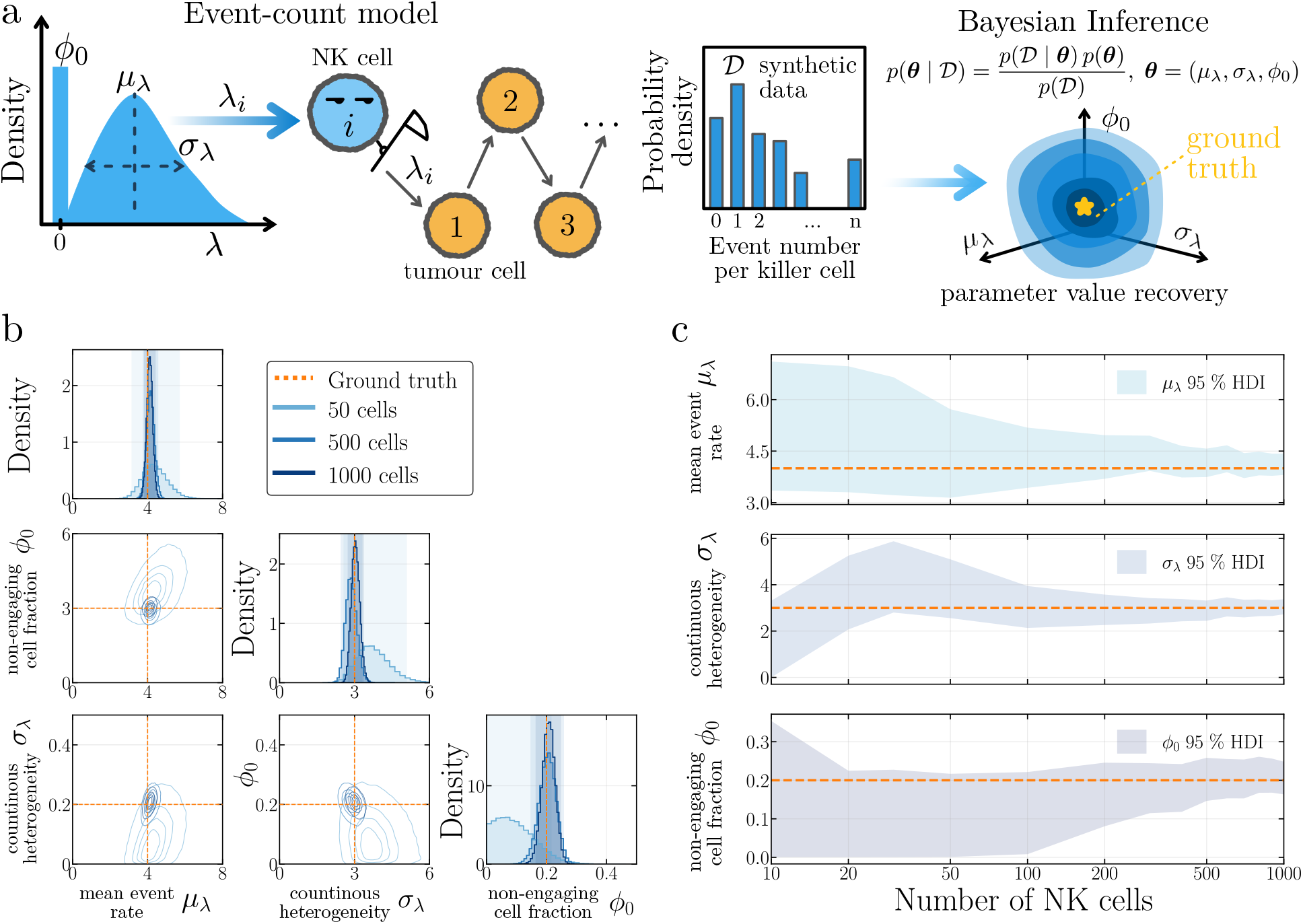
Validation of the Bayesian inference framework for NK cell interaction heterogeneity. **(a)** Schematic of the event count model and synthetic validation workflow. **(b)** Posterior recovery of *µ*_*λ*_, *σ*_*λ*_, and *φ*_0_ for synthetic datasets containing 50, 500, or 1000 NK cells. Upper diagonal panels show marginal posterior distributions, and off diagonal panels show contour plots of the joint posterior densities. Orange dashed lines indicate the ground truth values used for simulation, *µ*_*λ*_ = 4, *σ*_*λ*_ = 3, and *φ*_0_ = 0.2. Shaded regions indicate the 95% highest density intervals. **(c)** Effect of sample size on posterior uncertainty. Shaded regions indicate the 95% highest density intervals for *µ*_*λ*_, *σ*_*λ*_, and *φ*_0_ across increasing numbers of NK cells. Orange dashed lines indicate the corresponding ground truth values (*µ*_*λ*_ = 4, *σ*_*λ*_ = 3, and *φ*_0_ = 0.2). The x axis is shown on a logarithmic scale.

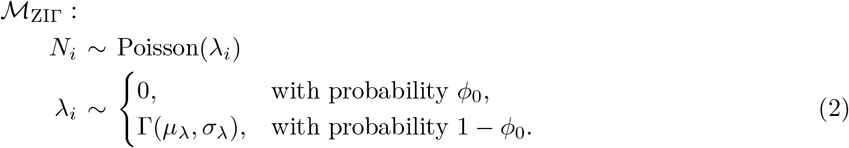

where *φ*_0_ denotes the fraction of non-engaging cells, and the Gamma distribution is parametrized by *µ*_*λ*_ and *σ*_*λ*_ describing the mean and standard deviation of event rates among the engaging population. The homogeneous Poisson model (eq. 1) emerges as a special case of this framework in the limit *φ*_0_ *→* 0 and *σ*_*λ*_*→* 0, corresponding to a population of statistically identical cells.

Before applying the framework to experimental data, we first tested whether the parameters describing the population structure could be reliably recovered from event-count distributions alone. To do so, we generated synthetic datasets under the hierarchical event-count model (Fig. 1a) using known parameter values and then performed Bayesian inference to infer the posterior distributions of the model parameters, allowing us to assess both the recovery of the ground truth parameters and the uncertainty associated with each estimate.

We repeated this analysis for increasing numbers of observed NK cells. As more single cell observations became available, the inferred parameter distributions became progressively narrower and centred around the ground truth values. In the parameter regime considered here, the posterior distributions became well constrained at approximately 500 cells, suggesting that the information contained within the event-count distribution is sufficient to robustly recover the underlying population structure (Fig. 1b,c).

Interestingly, different aspects of the population structure become identifiable at different sample sizes. For small numbers of cells (*≤* 100), the posterior distribution for the inactive fraction *φ*_0_ still included *φ*_0_ = 0, indicating that *<* 100 observed contact counts do not contain sufficient information to confidently infer the existence of a non-engaging subpopulation. In contrast, the posterior distribution for *σ*_*λ*_ remained clearly separated from zero even at low sample sizes, demonstrating that continuous cell-to-cell heterogeneity can be detected more readily than a discrete inactive subpopulation.

Having shown that the parameters of the heterogeneous model can be recovered from event-count distributions, we next asked whether the data contain sufficient information to distinguish between different origins of population heterogeneity. The homogeneous model (ℳ _homo_, eq. (1)) and the full zero-inflated Gamma model (ℳ_ZIΓ_ eq. (2)) represent the two extremes considered so far: a completely homogeneous population and a population containing both a non-engaging subpopulation and continuous cell-to-cell variability in event rates. To disentangle the contribution of these two sources of heterogeneity, we additionally considered two intermediate models. The first allows continuous heterogeneity but no non-engaging cells (labelled ℳ_Γ_, where *φ*_0_ = 0, *σ*_*λ*_ *≥* 0), while the second allows a non-engaging subpopulation but assumes identical event rates among the remaining cells (labelled ℳ_ZI_, where *φ*_0_ *≥* 0, *σ*_*λ*_ = 0), see Fig. 2a.

**Fig. 2.**
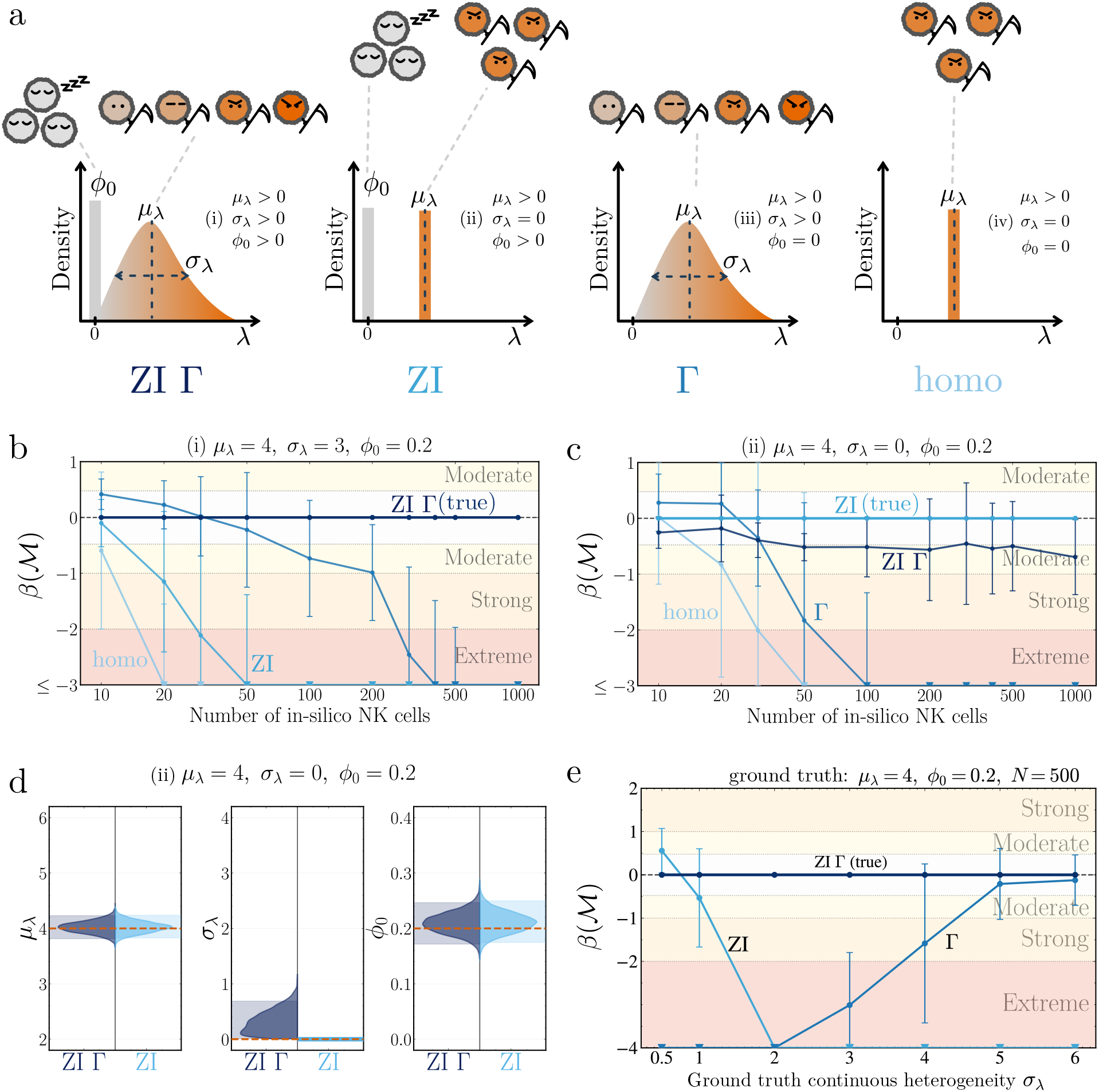
Validation of Bayesian model comparison for event-count population structures. **(a)** Schematic of the four candidate distributions for the cell-specific event rate *λ*_*i*_. **(b-c)** Model discrimination as a function of the number of cells for synthetic datasets generated under the ℳ_ZIΓ_ (left) and ℳ_Γ_ (right), respectively. Each synthetic dataset was analysed with all four candidate models. Model discrimination is expressed as the logarithm of the Bayes factor *β*(*M*) = log_10_ BF(*ℳ / ℳ* _true_), so the true model lies at zero and negative values indicate evidence against a candidate model relative to the true model. Points show the mean across independent synthetic replicates, and error bars show variability across 5 replicates. Background shading indicates conventional evidence ranges. **(d)** Posterior parameter distributions inferred from synthetic data generated under ℳ_ZI_, using the true model ℳ_ZI_ and the second most strongly supported model ℳ_ZIΓ_. Split violins show the marginal posterior distributions of *µ*_*λ*_, *σ*_*λ*_, and *φ*_0_. Orange dashed lines denote the ground truth values, and shaded regions denote 95% highest density intervals. **(e)** Sensitivity of model discrimination to the magnitude of continuous heterogeneity. Data were generated from ℳ_ZIΓ_ at *N* = 500, with *µ*_*λ*_ = 4 and *φ*_0_ = 0.2, while the ground truth *σ*_*λ*_ was varied.

A central challenge in this comparison is that increasingly heterogeneous models contain additional parameters and therefore have greater flexibility to reproduce the observed event-count distributions. Consequently, a better fit alone does not imply that the additional biological complexity is supported by the data. We therefore used Bayesian model comparison to determine whether the evidence contained within the event-count distributions justifies the inclusion of additional heterogeneity layers. Synthetic datasets were generated under each of the four population structures and analysed using all candidate models. For each dataset, we calculated the marginal likelihood of each model and compared them using Bayes factors, which quantify the relative support for competing biological hypotheses while accounting for differences in model complexity.

We report Bayes factors (BF) on a base-10 logarithmic scale relative to the true data-generating model, *β*(*M*) = log_10_ BF(*ℳ / ℳ*_true_). Values of *β* indicate equal support for the candidate and true models, while increasingly negative values indicate evidence against the candidate model. Values below -1, -2, and -3 correspond to moderate, strong, and extreme evidence against the candidate model, respectively [22].

Across all four synthetic population structures, the framework successfully recovered the correct origin of heterogeneity from event-count distributions. For populations containing continuous heterogeneity (ℳ_Γ_ and ℳ _ZIΓ_), the true model was clearly distinguished from the alternatives once approximately 200 cells were observed, with Bayes factors reaching strong to extreme evidence in favour of the correct population structure (Fig. 2b & Fig. S1a).

The most challenging scenarios corresponded to the homogeneous (ℳ _homo_) and zero-inflated homogeneous (ℳ _ZI_) populations (Fig. 2c & Fig. S1b). In these cases, the true models are limiting cases of the more flexible heterogeneous models, making it intrinsically difficult to rule out the presence of a weakly heterogeneous active population. Consequently, even with 1000 observed cells, the evidence against continuous heterogeneity remained moderate. However, inspection of the posterior distributions revealed that the inferred heterogeneity parameter concentrated near *σ*_*λ*_ = 0, while retaining support for the appropriate value for the rest of the parameters (Fig. 2d & Fig. S1c). Thus, although Bayesian model comparison alone provided only moderate separation, the inferred population structure remained fully consistent with the ground truth mechanism.

To further determine when population heterogeneity can be reliably inferred, we varied the continuous heterogeneity, *σ*_*λ*_, and the proportion of non-engaging cells, *φ*_0_, in datasets of 500 cells (Fig. 2e). At intermediate values of *σ*_*λ*_, the framework correctly identified ℳ _ZIΓ_. Discrimination weakened only at the extremes. When *σ*_*λ*_ ≪ *µ*_*λ*_, continuous heterogeneity becomes negligible and ℳ_ZIΓ_ approaches ℳ_ZI_. Conversely, when *σ*_*λ*_≫ *µ*_*λ*_, the broad Gamma distribution includes many rates close to zero, producing many cells with no events without requiring a distinct non-engaging population. As a result, ℳ_ZIΓ_ becomes difficult to distinguish from ℳ_Γ_ (Fig. 2e). The same principle applies to *φ*_0_: when *φ*_0_ is small, ℳ_ZIΓ_ approaches ℳ_Γ_, whereas evidence for a separate non-engaging population strengthens as *φ*_0_ increases (Fig. S1d). Thus, model discrimination is reliable across intermediate parameter regimes and weakens only when competing population structures become effectively identical.

### 2.3 Contact and killing distributions reveal distinct heterogeneity mechanisms across antibody treatments

Having established that the framework can reliably recover population structure from synthetic datasets, we next applied it to experimental time-lapse imaging data of NK cell cytotoxicity. We analysed untreated NK cells together with cells treated with rituximab or bispecific CC-96673. For each condition, we extracted two complementary summaries from the contact-kill trajectories: the total number of target encounters performed by each NK cell (*N*_*i*_) and the total number of successful killing events (*K*_*i*_). Contact counts quantify heterogeneity in target engagement, whereas kill counts quantify heterogeneity in realised cytotoxic output. Each condition contained approximately 500 analysed NK cells, placing the data within the regime where the synthetic analyses indicated reliable parameter and model recovery. Interestingly, the overall event-count distributions appeared broadly similar across both treatments (Fig. 3b), making them an ideal test case for determining whether mechanistic differences can be inferred from seemingly similar population-level behaviours.

**Fig. 3.**
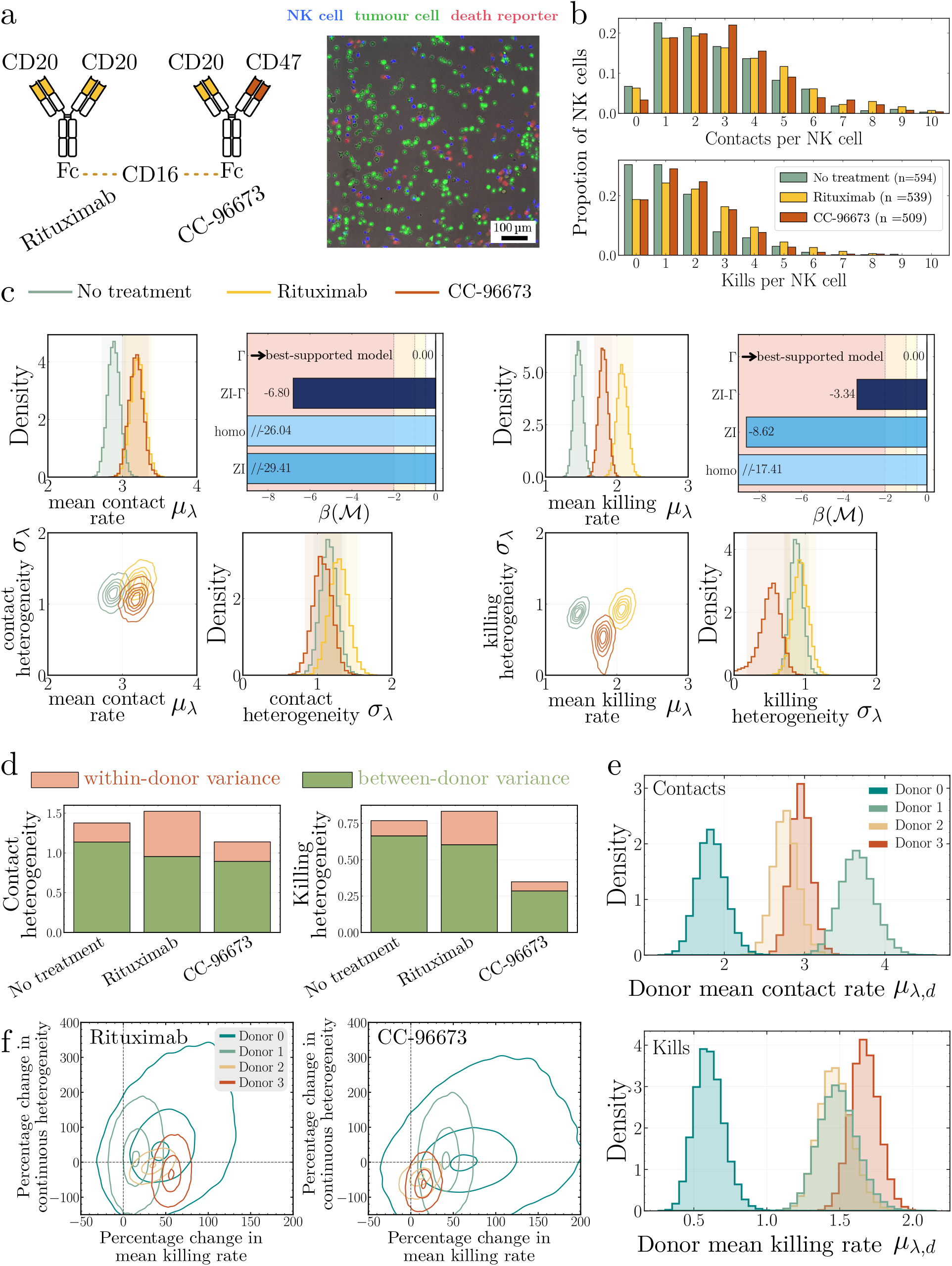
Event-count analysis of NK contact and killing distributions for different antibody treatments. **(a)** Schematic of rituximab and CC-96673 treatments, with a representative time-lapse imaging field. **(b)** Empirical distributions of target contacts and kills per NK cell under no treatment, rituximab, and CC-96673 treatment. The number of analysed NK cells is shown in the legend. **(c)** Bayesian inference of the mean event rate *µ*_*λ*_ and population heterogeneity *σ*_*λ*_ for contact (left) and kill (right) counts. Marginal and joint posterior distributions are shown for each treatment condition. Insets show Bayesian comparison of ℳ_homo_, ℳ_ZI_, ℳ_Γ_, and ℳ_ZIΓ_. Bayes factors are reported relative to the best-supported model. **(d)** Decomposition of the population continuous variance expectation into within-donor (salmon) and between-donor (green) components for contact counts (left) and kill counts (right). **(e)** Marginal posterior distributions of the donor specific mean event rate *µ*_*λ,d*_ under the no-treatment condition for contact counts (top) and kill counts (bottom). **(f)** Joint posterior distributions of donor specific percentage changes in the mean killing rate *µ*_*λ,d*_ and continuous heterogeneity *σ*_*λ,d*_, relative to the no treatment condition. Contrasts are shown for rituximab (left) and CC-96673 treatment (right). Contours outline the 5%, 50%, and 95% highest density regions of the bivariate posterior distribution.

We first asked which heterogeneity structure best explains the observed contact and killing distributions. To address this, we compared the four models introduced above, representing different combinations of continuous heterogeneity and non-engaging subpopulations. For both contact counts and killing counts, Bayesian model comparison consistently favoured the ℳ_Γ_ model across all experimental conditions (Fig. 3c insets). This indicates that both target engagement and cytotoxic activity are best described by a continuum of cellular behaviours spanning a broad range of interaction and killing propensities, rather than a homogeneous model. Conversely, we found no evidence supporting the existence of a distinct non-engaging subpopulation, either at the level of target encounters or successful killing events. Importantly, this conclusion was consistent across untreated, rituximab-treated, and bispecific-antibody-treated populations, suggesting that continuous cell-to-cell heterogeneity is a robust feature of NK cell behaviour under all conditions examined. This conclusion was further supported by inference under the most flexible model,ℳ_ZIΓ_. Across all conditions, the posterior distribution of the non-engaging fraction, *φ*_0_, concentrated near zero (Fig. S2a). For kill counts under rituximab and CC-96673 treatment, the Bayes factor separation between ℳ_Γ_ and the alternative heterogeneous models was relatively modest (Fig. S2b). However, the posterior distributions under ℳ_ZIΓ_ led to the same biological interpretation, with substantial continuous heterogeneity but negligible support for zero inflation.

Having established that all conditions are best described by continuous heterogeneity, we next compared the inferred contact-rate distributions across treatments. The posterior distributions of the heterogeneity parameter, *σ*_*λ*_, largely overlapped across untreated, rituximab-treated, and bispecific-antibody-treated populations, indicating similar levels of cell-to-cell variability in target engagement (Fig. 3c left). In contrast, both antibody treatments shifted the posterior distribution of the mean contact rate, *µ*_*λ*_, towards higher values relative to untreated cells. These results suggest that antibody treatment increases the frequency of target sustained encounters while leaving the degree of heterogeneity in contact behaviour largely unchanged.

A different picture emerged when analysing the killing-rate distributions (Fig. 3c right). Although both rituximab and the bispecific CC-96673 increased cytotoxic activity relative to untreated cells, they did so through distinct changes in the inferred population structure. Rituximab produced the largest increase in the mean killing rate, *µ*_*λ*_, indicating an overall enhancement of cytotoxic efficiency across the NK cell population. In contrast, the CC-96673 produced a more modest increase in *µ*_*λ*_ but substantially reduced the heterogeneity parameter, *σ*_*λ*_. Thus, rather than primarily increasing the average killing activity, the CC-96673 appeared to make cytotoxic responses more uniform across the population. These results demonstrate that similar population-level killing profiles can arise from fundamentally different underlying mechanisms: one driven by an increase in killing efficiency, and the other by a reduction in cell-to-cell variability.

We then asked whether the heterogeneity in NK cell behaviour arises from the cell-to-cell variation within each donor or from differences between donors. The event-count analysis above treated NK cells from all donors as a single population; therefore we extended the same four event-count models using a donor-aware hierarchical framework (methods 4.6). This donor-aware framework allows the mean event rate, continuous cell-to-cell heterogeneity, and non-engaging cell fraction to vary between donors.

Through the model comparison between the donor-aware extension of all four models (ℳ_homo_, ℳ_ZI_, ℳ_Γ_, and ℳ_ZIΓ_), the ℳ_Γ_ remained the best-supported model for both contact and kill counts (Fig. S3b). Moreover, the population parameters reconstructed from the donor-specific posteriors closely agreed with those obtained from the analysis that did not distinguish donors (Fig. 3c & Fig. S3a). Thus, the donor-aware analysis did not alter the conclusions mentioned above, but allowed us to investigate the source of heterogeneity observed at NK cell population level in a more detailed perspective.

In particular, variance decomposition showed that population-level cytotoxic variability arises from both cell-to-cell heterogeneity within donors and systematic differences between donors, with the latter accounting for a larger share (Fig. 3d). Analysis of the donor-specific mean contact and killing rates reveals this pre-existing heterogeneity even before treatment (Fig. 3e and S5).

Given the substantial between-donor heterogeneity observed, we next asked whether the effects of antibody treatment on NK cell killing behaviour varied among donors and, specifically, whether some donors exhibited greater treatment responsiveness than others. Rituximab increased the mean kill rate in donors 2 and 3 by approximately 40% and 60%, respectively (Fig. 3f, left). By contrast, donor 1 showed the clearest response to the CC-96673, with an approximately 40% increase in mean kill rate relative to no treatment (Fig. 3f, right). Thus, the two antibody treatments differed in which donors exhibited the strongest increase in NK cell killing activity. Additionally, CC-96673 treatment exhibits a tendency of reducing the within-donor continuous heterogeneity in donor 2 and donor 3 as reported in the population trends, with rituximab primarily increasing the mean kill rate whereas the CC-96673 reduced continuous heterogeneity.

Finally, we also asked whether kill-count heterogeneity could be explained by a distinct subpopulation of cells with elevated killing rates (Methods 4.7). Across untreated, rituximab and CC-96673 conditions, the data did not support this additional component (Fig. S7), providing no evidence for a separate higher-rate “superkiller” group within donors under the assumptions and priors considered.

### 2.4 A trajectory model separates cellular heterogeneity from history dependent killing

The event-count analysis revealed substantial heterogeneity in both target engagement and cytotoxic output across NK cell populations. However, it remains unclear whether this variability reflects stable intrinsic differences between cells or emerges through the accumulation of previous interactions. For example, cells that perform many killing events may represent a pre-existing highly cytotoxic subpopulation. Alternatively, such heterogeneity could emerge dynamically if previous successful or unsuccessful encounters modify the probability of future killing events. The summary statistics *N*_*i*_ and *K*_*i*_ cannot distinguish between these possibilities because they record only how many contacts and killing events occurred, not the order in which they took place.

To address this limitation, we extended the framework to incorporate the full interaction history of each cell. Rather than summarising the number of successful kills (*K*_*i*_), we considered the ordered sequence of contact outcomes, *x*_*i*_ = (0, 0, 1, …), where each element records whether a particular target encounter was lethal or non-lethal. Unlike event counts, these trajectories retain information about the temporal arrangement of killing events. For example, a cell that kills during its first three encounters and then ceases killing contains a very different biological signature from a cell that kills only after several unsuccessful contacts, even if both cells perform the same total number of kills. The ordered trajectories therefore provide a means to test whether previous interactions influence future cytotoxic decisions.

In order to incorporate this history effect, we model the outcome of each target encounter as a binary cytotoxic decision whose probability of success depends on both the intrinsic killing propensity of the cell and its previous interaction history. Specifically,

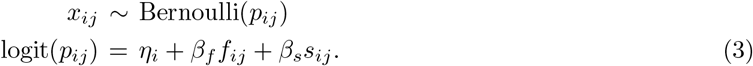

Here, *η*_*i*_ represents the baseline killing propensity of cell *i*, while *f*_*ij*_ and *s*_*ij*_ denote the numbers of previous failed and successful killing events experienced before contact *j*, respectively. The coefficients *β*_*f*_ and *β*_*s*_ quantify how previous encounters influence future killing decisions. Positive values correspond to cytotoxic activation, whereby previous encounters increase the probability of subsequent killing, whereas negative values correspond to inhibition or exhaustion-like behaviour.

As in the event-count framework, baseline killing propensities were allowed to vary across the population to capture history-independent cell-to-cell heterogeneity,

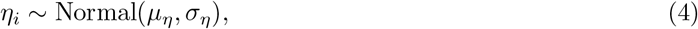

where *µ*_*η*_ describes the average baseline killing propensity of the population and *σ*_*η*_ quantifies cell-to-cell heterogeneity. In this formulation, *σ*_*η*_ captures stable differences between NK cells, whereas *β*_*f*_ and *β*_*s*_ capture history-dependent changes in killing probability induced by previous encounters.

We validated whether the full trajectory model could recover known parameters from synthetic contact kill histories. For different synthetic datasets, we examined posterior recovery as the number of observed NK cells increased. At small sample size, posterior uncertainty was broad, particularly for the decision-making parameters *µ*_*η*_, *σ*_*η*_, *β*_*f*_ , and *β*_*s*_ (Fig. 4b). This is expected because these parameters are informed not only by the total number of cells, but also by the number and order of contacts within each single cell trajectory. In contrast, the contact rate parameters *µ*_*λ*_ and *σ*_*λ*_ were more directly informed by the per cell contact counts and were therefore recovered more rapidly. As the number of in silico NK cells increased, posterior intervals narrowed and posterior mass concentrated around the ground truth values (Fig. 4b). By approximately 500 cells, the model recovered the main features of the simulated process, including the mean contact rate, the degree of contact heterogeneity, the population mean baseline killing propensity, and the signs of both history dependent effects.

**Fig. 4.**
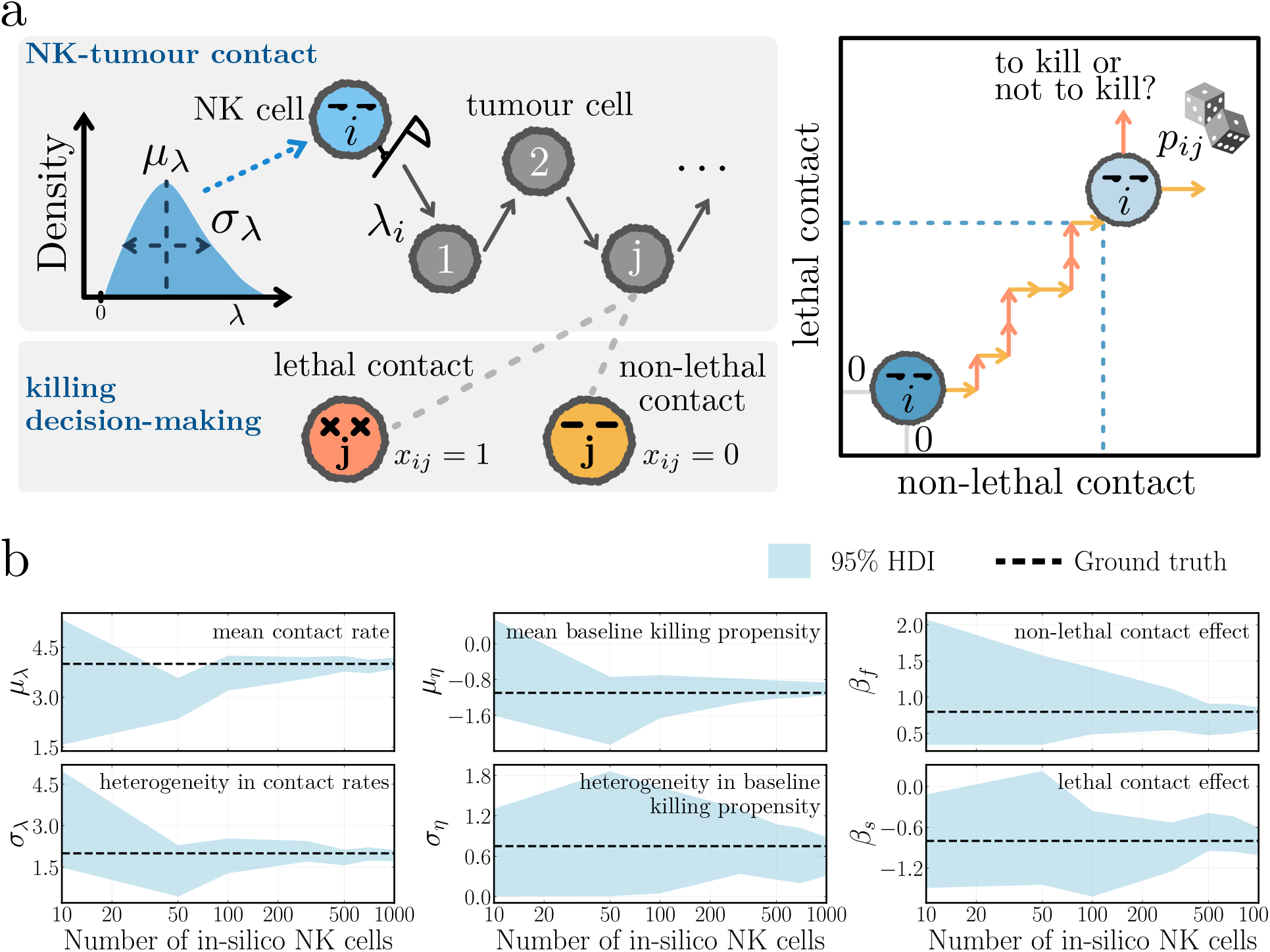
Validation of the full contact-kill trajectory model using synthetic data. **(a)** Schematic of the trajectory model. Contact formation is represented by a Gamma distributed cell specific contact rate *λ*_*i*_, with population mean *µ*_*λ*_ and standard deviation *σ*_*λ*_. Conditional on contact formation, each contact outcome is modelled as a Bernoulli killing decision. The probability of killing depends on the baseline killing propensity *η*_*i*_, the number of previous non lethal contacts *x*_*ij*_ , and the number of previous lethal contacts *y*_*ij*_. **(b)** Effect of sample size on posterior uncertainty for the full model. Shaded regions indicate the 95% highest density intervals for *µ*_*λ*_, *σ*_*λ*_, *µ*_*η*_, *σ*_*η*_, *β*_*f*_ , and *β*_*s*_ across increasing numbers of in silico NK cells. Orange dashed lines indicate the ground truth values used for simulation.

### 2.5 Trajectory models reveal treatment dependent cytotoxic decision-making

We next applied the validated trajectory framework to imaging-derived contact-kill histories from the same experimental antibody conditions presented in Section 2.3. Analysis of lethal and non-lethal contacts shows variability in behaviour depending on the history of each cell (Fig. 5a). At the origin, where no previous contacts had occurred, the empirical probability that the first contact was lethal was slightly higher under both antibody treatments than under no treatment. The fraction of cells reaching each state decreased rapidly as the number of accumulated contacts increased, resulting in noisier empirical estimates for sparsely populated states. The state maps also suggested an apparent separation of trajectories: some cells underwent repeated lethal contacts, whereas others accumulated predominantly non-lethal contacts. Accordingly, cells with more previous kills appeared more likely to kill again, while cells with more previous non-lethal contacts appeared less likely to kill subsequently.

**Fig. 5.**
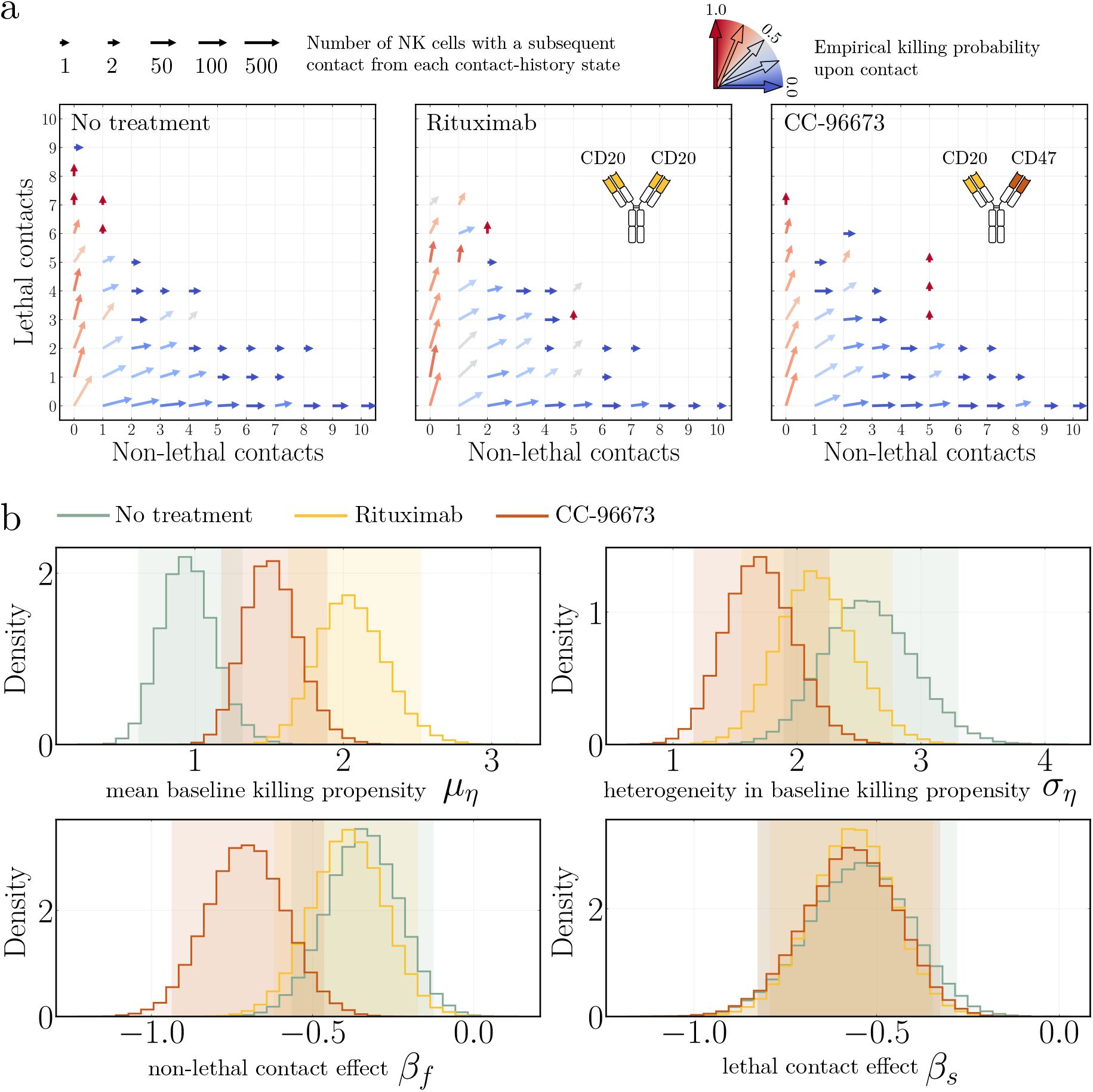
Trajectory analysis reveals heterogeneous and history-dependent NK cell cytotoxic decisions. **(a)** Empirical contact-history state maps for no treatment, rituximab, and CC-96673 treatment. Arrow direction and colour both encode the empirical probability that the next contact is lethal at that history state, whereas arrow-tail length is proportional to the number of observed transitions originated from each history state. **(b)** Marginal posterior distributions under the heterogeneous history-dependent model. Shown are the population mean baseline killing propensity (*µ*_*η*_), cell-to-cell heterogeneity in baseline killing propensity (*σ*_*η*_), and the effects of previous non-lethal (*β*_*f*_) and lethal (*β*_*s*_) contacts on the probability that a subsequent contact is lethal. Coloured shaded areas indicate 95% highest density intervals.

The result of the Bayesian analysis supports the conclusion that cell heterogeneity is a combination of stable cell-to-cell differences in baseline killing propensity and changes in killing behaviour induced by previous contacts; neither mechanism alone was sufficient to describe the data (Fig. S6). The inferred effects on baseline killing propensity were consistent with the event-count analysis. Both treatments increased the mean baseline killing propensity *µ*_*η*_, with a more pronounced increase under rituximab (Fig. 5b). In addition, rituximab had little effect on cell-to-cell heterogeneity *σ*_*η*_, whereas the CC-96673 reduced this variability, resulting in more consistent cytotoxic behaviour across the NK cell population.

Analysis of the history-dependent coefficients supported an overall decline in the killing rate for all three experimental conditions after lethal or non-lethal contacts *β*_*s*_, *β*_*f*_ *<* 0 (Fig. 5b). The distributions for the effect of lethal contacts *β*_*s*_ overlapped substantially across the three conditions, providing little evidence that either antibody altered the effect of previous successful kills. By contrast, analysis of the effect of non-lethal contacts *β*_*f*_ showed a stronger effect for the CC-96673 than under no treatment and rituximab, respectively; suggesting that the CC-96673 lose their killing capability at faster rate after non-lethal contacts than rituximab or untreated cells.

### 2.6 BARRACUDA provides a reusable implementation of the framework

To make the analyses developed here accessible for other datasets, we created BARRACUDA (Bayesian Analysis of Randomness, Resolving Alternative Causes of Uncertainty in Differential Activity), an open-source platform for analysing single-cell cytotoxicity. The web application combines event-count and ordered contact-history analyses within a common workflow. Users can compare alternative population models for contact or kill counts, separate within- and between-donor variation, and distinguish stable differences in killing propensity from changes associated with previous encounters.

The platform also includes synthetic examples, downloadable outputs, and example notebooks. The accompanying cyto-barracuda Python package provides programmatic access to the same simulation and inference routines for reproducible scripted analyses. Together, these interfaces provide a directly usable implementation of the framework developed in this study.

## 3 Discussion

In this study, we established two Bayesian inference frameworks for investigating the origins of heterogeneity in NK cell cytotoxicity. The first analyses the numbers of target cell contacts and kills recorded for individual NK cells. It distinguishes stochastic variation from different origins of population heterogeneity: whether the data observed can be explained by chance alone, continuous cell-to-cell differences in contact or killing rates, a distinct non-engaging subpopulation, or a combination of these mechanisms. By incorporating donor identity, this framework further separates variation within and between donors, enabling us to quantify how donor-to-donor differences contribute to the heterogeneity observed at the population level. The second, trajectory-based framework retains the order of lethal and non-lethal contacts for each NK cell, allowing us to distinguish the stable intrinsic killing propensities of individual cells from changes associated with their previous interactions (intrinsic heterogeneity vs longitudinal heterogeneity). Together, these frameworks connect observable single-cell behaviours to hidden biological processes and demonstrate how increasingly detailed trajectory information can answer progressively more specific questions about the mechanistic origins of NK cell decision-making.

When applied to experimental data, the analysis showed how timelapse interaction data contain plenty of mechanistic information on the sources of heterogeneity. We found no evidence for a distinct subpopulation of cells that was intrinsically unable to contact or kill targets during the assay. Instead, NK cells occupied a continuum ranging from cells with low interaction or killing activity to highly active cells. This distinction is biologically important because cells making no contacts or kills during a finite imaging period might otherwise be labelled inactive, whereas our framework shows that they do not constitute a stable inactive population. Such cells may instead occupy the lower end of a continuous activity distribution or may simply have experienced no events because of stochasticity. This highlights how behavioural classifications such as non-killing or killing subsets may be insufficient [9], as they do not necessarily correspond to discrete and stable cellular identities.

The event-count analysis suggested that the two antibody treatments reshaped NK cell behaviour through distinct mechanisms. Both increased the average number of target contacts. But their effects on killing differed: rituximab primarily increased the mean killing rate, whereas the CC-96673 produced a more modest increase in mean activity but markedly reduced cell-to-cell variability. Thus, treatments producing similar overall numbers of killing events can generate different distributions of cellular behaviour. Increasing the mean enhances population-wide activity, whereas reducing heterogeneity limits the proportion of weakly responsive cells and produces a more consistent cytotoxic response. Distinguishing effects on mean activity from effects on heterogeneity also points to potentially distinct molecular mechanisms underlying the two treatments. These differences may be related to how the antibodies are organised on the target cell surface. Rituximab increased the proportion of target cells exhibiting CD20 capping, whereas little capping was observed with CC-96673 or no treatment (Fig. S8c). One previous study showed that rituximab-induced target cell capping increased NK cell-mediated cytotoxicity, especially when the contact happened at the capping site [23]. Therefore one speculative explanation would be that allowing us to distinguish rituximab-induce capping generates strong but spatially restricted Fc-mediated stimulation, whereas dual CD20/CD47 engagement by CC-96673 may provide a more consistent signal across target cell encounters. Quantifying both therefore provides a fuller picture of how treatment reshapes NK cell function. Extending this analysis across donors showed that treatments can reshape NK cell function differently between individuals. By resolving these donor-specific responses, the framework could help identify which interventions are most effective for each donor and ultimately support more personalised therapeutic strategies.

Longitudinal analysis allowed us to separate differences in killing propensity from changes induced by previous encounters. Previous lethal contacts reduced subsequent killing similarly across conditions. However, a suppressive effect of non-lethal contacts was stronger when cells were incubated with the CC-96673. Thus, the antibodies differed not only in how they altered intrinsic cytotoxicity, but also in how NK cells responded according to their interaction history. This distinction opens the door to determining which molecular processes, including altered receptor signalling, immune synapse formation, degranulation, or cytotoxic granule availability, underlie the different effects of successive encounters.

An advantage of the Bayesian framework is that it reveals not only which behavioural effects are supported by the data, but also where the available experiments are insufficient. Broad posterior distributions identify poorly constrained quantities and therefore indicate where additional measurements would be most informative. This was particularly apparent for several donor-level parameters, reflecting the limited number of donors examined. The purpose of the present study was not to establish definitive donor-specific responses to rituximab or the CC-96673, but to show how such differences can be quantified and used to direct subsequent experiments. Larger donor cohorts will therefore be required to determine which of these effects are reproducible and biologically meaningful.

The models also make simplifying assumptions about population structure, including that cellular propensities follow a unimodal continuous distribution. More complex populations may contain discrete subgroups, including rare highly cytotoxic or super-killer cells, and could instead be represented using mixture distributions. Comparing such models would help determine whether extreme serial-killing behaviour reflects the tail of a continuous distribution or a distinct cellular state.

Although the framework identifies behavioural differences, it does not determine the molecular processes that generate them. A more mechanistic extension would couple the trajectory analysis to agent-based models describing processes such as receptor signalling, immune-synapse formation, degranulation, and cytotoxic-granule availability. Bayesian inference for such detailed and computationally expensive models would likely require simulation-based approaches, including neural posterior estimators [24]. Combining these models with transcriptomic, proteomic, or phenotypic measurements from functionally characterised cells could then connect inferred behavioural mechanisms to molecular state.

Finally, extracting contact and killing histories from time-lapse imaging remains labour intensive and depends on the accuracy of cell segmentation, tracking, and event-classification tools. This restricts the number of cells, donors, and treatment conditions that can be analysed. Reducing each trajectory to ordered contact and killing events also simplifies the underlying interactions by omitting contact duration and spatial information, including the number and arrangement of target cells contacting an NK cell simultaneously. Incorporating this additional spatiotemporal resolution could reveal how local cellular context shapes cytotoxic decisions. Automated microscopy and analysis pipelines will be required to increase throughput, while microwell systems combining semi-automated functional screening with retrieval of selected NK cells provide one possible route towards linking cytotoxic behaviour with downstream molecular analysis [10].

## 4 Methods

### 4.1 Primary NK cell isolation and culture

Blood leucocyte cones were obtained from the National Blood Transfusion Service in Manchester and London, UK, in accordance with ethical guidelines (Manchester License: 05/Q0401/108 and 22/SW/0076 for Imperial College London). PBMCs from healthy adult donors were isolated by density centrifugation using a Ficoll gradient. NK cells were isolated using a negative magnetic selection kit (Miltenyi Biotec). Cells were cultured at 1 *×* 10^6^ cells/mL in clone medium consisting of Dulbecco’s Modified Eagle Medium (DMEM) supplemented with 30% Ham’s F12, 10% human serum, 1% non-essential amino acids, 2 mM L-glutamine, 50 U/mL penicillin–streptomycin, 50 µM 2-mercaptoethanol, and 1 mM sodium pyruvate. Unless otherwise stated, reagents were obtained from Sigma–Aldrich, except for 2-mercaptoethanol and sodium pyruvate, which were obtained from Gibco. Clone medium was supplemented with 200 U/mL recombinant IL-2 (Gibco) immediately after isolation. NK cells were rested at 37 ^*°*^C and 5% CO_2_, and used 6 or 7 days after isolation.

### 4.2 Cell lines

The 721.221 cell lines were from Shimizu and DeMars. The cells were cultured at 37°C and 5% CO_2_ in RPMI 1640 medium supplemented with 10% FBS, 1% non-essential amino acids, 1 mM sodium pyruvate, 2 mM L-glutamine, and 50 U/mL penicillin/streptomycin.

### 4.3 Antibodies

The bispecific antibody used in this study was CC-96673 (BMS-986358), a humanised CD20 *×* CD47 IgG1 antibody with a high affinity CD20 binding arm, an affinity detuned CD47 binding arm, and a wild type IgG1 Fc region [20]. Rituximab, a monospecific bivalent antibody targeting CD20, was used as the comparator.

### 4.4 Time lapse imaging killing assay

Human NK cells and 721.221 target cells were labelled separately with CellTrace^TM^ calcein red-orange and calcein green AM (Thermo Fisher Scientific), respectively. Each dye was used at a concentration of 1 µg/mL in PBS, and cells were incubated for 20 min at 37 ^*°*^C. Following labelling, cells were washed with serum containing RPMI 1640 medium and resuspended in fresh complete RPMI 1640 medium. The NK cell and target cell suspensions were adjusted to 84,000 and 336,000 cells/mL, respectively.

Wells of an ibidi µ-Slide 18 Well chamber slide were coated with fibronectin in PBS and washed with PBS before cell addition. For antibody treated conditions, antibody was added to the target cell suspension before the NK cells were introduced. The amount of antibody was adjusted to give a final well concentration of 100 ng/mL. Equal volumes of the NK cell and target cell suspensions were then combined, producing an effector to target ratio of 1:4.

Immediately after mixing, 90 µL of the combined cell suspension was added to each well. This corresponded to approximately 3,780 NK cells and 15,120 target cells per well. The chamber slide was centrifuged at 50 *× g* for 1 min to bring the suspended cells to the bottom of each well. Subsequently, 10 µL of TO-PRO-3 iodide solution was added, giving a final well volume of 100 µL and a final TO-PRO-3 dilution of 1:1000.

Cells were imaged immediately using an inverted Nikon Ti Eclipse microscope in widefield mode with a 10 *×* objective. The chamber was maintained at 37 ^*°*^C and 5% CO_2_ during image acquisition. Images were acquired every 3 min for 5 h.

Image sequences were analysed manually. Contacts were identified as instances physical contact between an NK cell and a tumour cell lasting for at least three consecutive frames, while tumour cell death was identified by TO-PRO-3 fluorescence.

### 4.5 Event count models, likelihoods, and priors

We used Bayesian event count models to analyse single cell interaction histories. An event denotes either a target contact or a target kill, depending on the analysis. For each NK cell *i* = 1, … , *M* , the observed event count over an imaging window of duration *T* was denoted by *N*_*i*_. The data were written as **N** = (*N*_1_, … , *N*_*M*_). The same framework was applied separately to contact counts and kill counts.

We compared four candidate models: a homogeneous Poisson model ℳ_homo_, a zero inflated Poisson model ℳ_ZI_, a heterogeneous Gamma Poisson model ℳ_Γ_, and a zero inflated heterogeneous Gamma Poisson model ℳ_ZIΓ_. These models differ in whether variation in event counts is attributed to Poisson sampling noise, structural inactivity, continuous cell heterogeneity, or a combination of these mechanisms.

In the homogeneous model, all cells shared a common event rate *λ*:

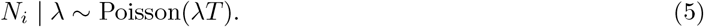

The corresponding probability mass function was

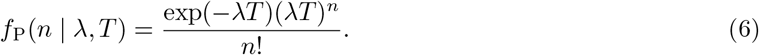

In the zero inflated Poisson model, cells were inactive with probability *p*_0_ and otherwise followed the homogeneous Poisson model:

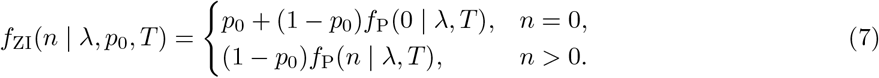

In the heterogeneous Gamma Poisson model, each cell had its own event rate,

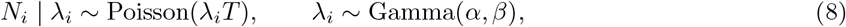

where *α* is the shape parameter and *β* is the rate parameter. We used the mean and standard deviation parameterisation

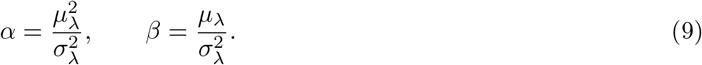

The latent rates *λ*_*i*_ were analytically marginalised. This gives the Gamma Poisson probability mass function

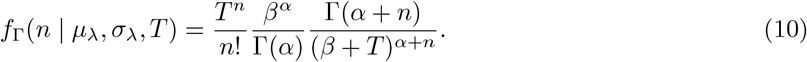

For compact notation, we write

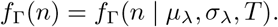

In the zero inflated heterogeneous Gamma Poisson model, cells were inactive with probability *p*_0_, while active cells followed the Gamma Poisson model:

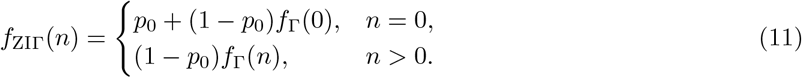

For each model ℳ with parameter vector *θ*_*M*_, the likelihood was

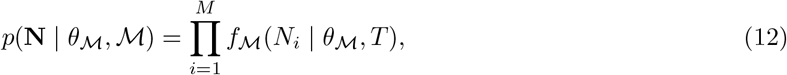

and the log likelihood was

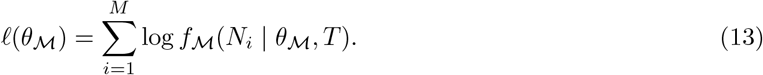

Unless otherwise stated, event rates were assigned priors on a base 10 logarithmic scale,

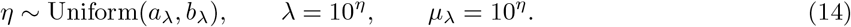

The inactive fraction and heterogeneity scale were assigned

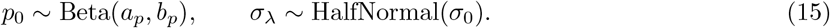

We used *a*_*λ*_ =*™*5, *b*_*λ*_ = 2, *a*_*p*_ = 1, *b*_*p*_ = 1, and *σ*_0_ = 1 unless otherwise stated.

Posterior inference was performed using Sequential Monte Carlo sampling in PyMC [25]. For the Gamma

Poisson and zero inflated Gamma Poisson models, the analytically marginalised log likelihoods were implemented directly as model potentials. Full count normalisation terms were retained for marginal likelihood estimation and Bayes factor comparison.

### 4.6 Donor-aware extensions of the event counts inference framework

To account for variation between donors, we constructed a hierarchical extension of the event-count models described above. We first introduced shared reference parameters for the mean event rate, continuous heterogeneity, and non-engaging fraction,

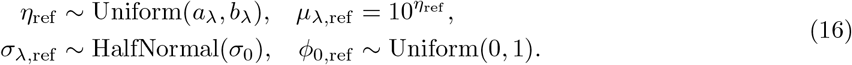

For each donor *d*, donor-specific parameters were generated by applying zero-centred deviations on unconstrained scales,

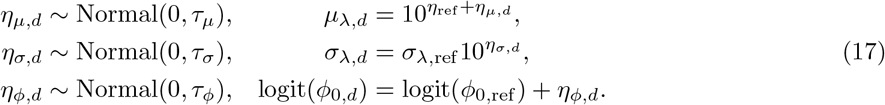

A deviation of zero corresponds to the shared reference value. The prior hyperparameters were fixed across conditions and outcomes as *a*_*λ*_ = *™*1.5, *b*_*λ*_ = 1.5, *σ*_0_ = 3, *τ*_*µ*_ = 0.3, *τ*_*σ*_ = 0.3, and *τ*_*φ*_ = 1. Priors involving *σ*_*λ,d*_ or *φ*_0,*d*_ were excluded from models that did not contain the corresponding source of heterogeneity.

The donor-specific parameters were inferred jointly. The population mean event rate 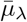, continuous heterogeneity 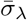, and non-engaging fraction 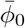 were not sampled as additional model parameters, but were reconstructed from the donor-specific posterior distributions. Let *w*_*d*_ denote the proportion of cells contributed by donor *d*, and define the donor weight among engaging cells as

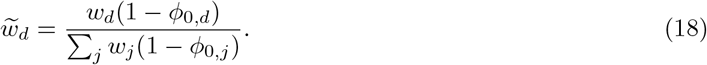

At each posterior draw, the population parameters were calculated as

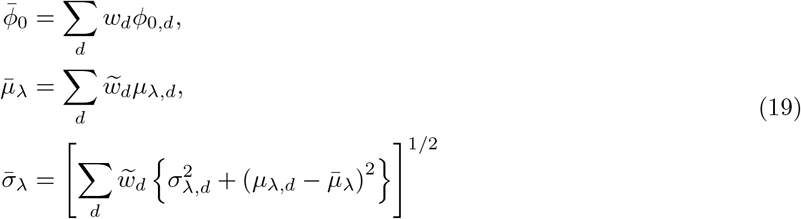

The population level continuous variance can be decomposed as

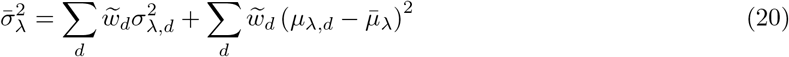

where the first term 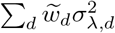 quantifies the weighted average of cell-to-cell variance within donors, whereas the second term 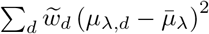 quantifies the weighted between-donor variance.

Posterior inference and marginal likelihood estimation used Sequential Monte Carlo sampling with 10,000 particles in each of 8 chains. Treatment effects were quantified using posterior contrasts. For treatments *A* and *B*,

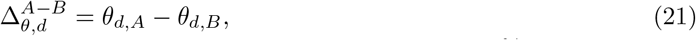

where *θ* denotes either *µ*_*λ*_ or *σ*_*λ*_. For each contrast, we report its posterior median, 95% highest density interval, and the posterior probability that it was positive or negative.

### 4.7 Donor-aware 2Γ model

We further extended the donor-aware Γ model to a 2Γ model to assess support for an additional higher-rate component. For each donor *d*, the first Γ component retained the mean event rate *µ*_*λ,d*_, continuous heterogeneity *σ*_*λ,d*_, and hierarchy and priors described above. The second component had fraction *π*_*d*_, mean event rate *r*_*d*_*µ*_*λ,d*_, with *r*_*d*_ *>* 1, and separately estimated continuous heterogeneity *σ*_*λ,H,d*_.

The additional parameters were assigned independent priors across donors and independent of the first component’s hierarchy,

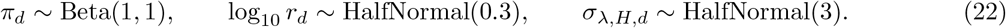

After analytically marginalising cell rates and component assignments, the count probability was

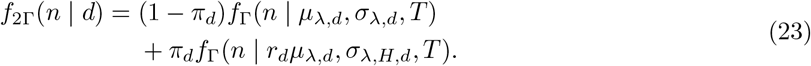

Posterior inference and marginal likelihood estimation followed the procedure above. Bayes factors compared the donor-aware Γ and 2Γ models jointly across donors, separately for contact and kill counts in each condition.

### 4.8 Bayes factors and marginal likelihoods

Model comparison was performed using Bayes factors. For each candidate model ℳ with parameters *θ* and data *D*, the marginal likelihood is defined as

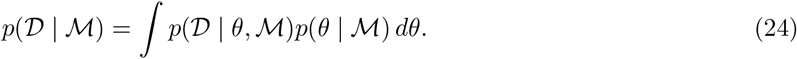

The marginal likelihood measures how well a model explains the data after averaging over its prior parameter space, and therefore accounts for both goodness of fit and model complexity [26].

Posterior inference and marginal likelihood estimation were performed using the Sequential Monte Carlo sampler implemented in PyMC [25]. The sampler uses a sequence of tempered distributions,

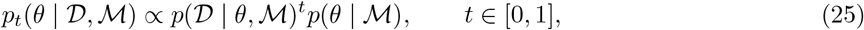

where *t* = 0 corresponds to the prior and *t* = 1 corresponds to the posterior. The log marginal likelihood can be estimated from this sequence using thermodynamic integration,

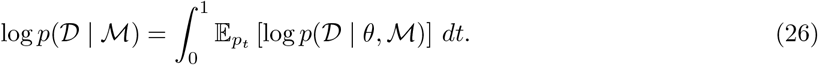

In practice, this integral was approximated numerically over the temperature ladder generated by the SMC sampler [27].

For two competing models, ℳ_1_ and ℳ_2_, model support was compared using the log Bayes factor,

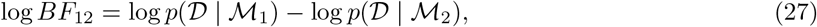

where *p*(*D*|ℳ) is the marginal likelihood of model ℳ. This is equivalent to taking the logarithm of the Bayes factor

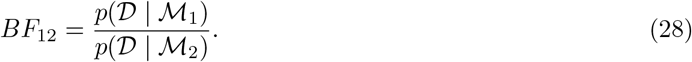

Values of log *BF*_12_ *>* 0 indicate support for ℳ_1_ over ℳ_2_, whereas values of log *BF*_12_ *<* 0 indicate support for ℳ_2_ over ℳ_1_.

## Data availability

Processed single-cell data are publicly available in the BARRACUDA repository at https://github.com/sthsci/Barracuda/tree/main/data. Raw time-lapse microscopy data are not publicly available. Enquiries regarding access to these data should be directed to Ruben Perez-Carrasco at.

## Code availability

The BARRACUDA source code and analysis scripts are publicly available under the MIT licence at https://github.com/sthsci/Barracuda. The Python package is distributed through PyPI at https://pypi.org/project/cyto-barracuda/, and the web application is accessible at https://barracuda.clingenland.science/. A software snapshot, version `v_zenodo_0.1.0`, has been deposited in Zenodo at https://doi.org/10.5281/zenodo.22085204; access to the archived files is currently restricted.

## Author contributions

Conceptualization: E.S., C.H., and R.P-C.; Methodology: E.S., C.H., and R.P-C.; Software development (BARRACUDA web application): E.S.; Formal analysis: E.S. and R.P-C.; Experimental: C.H. and K.S.H.; Resources: L.P. and L.S.; Data curation: E.S. and C.H.; Visualization: E.S. and C.H.; Writing, review and editing: E.S., C.H., D.M.D., and R.P-C.; Supervision: D.M.D. and R.P-C.; Funding acquisition: D.M.D. and R.P-C. All authors reviewed and approved the final manuscript.

## Supplementary Information

**Fig. S1.**
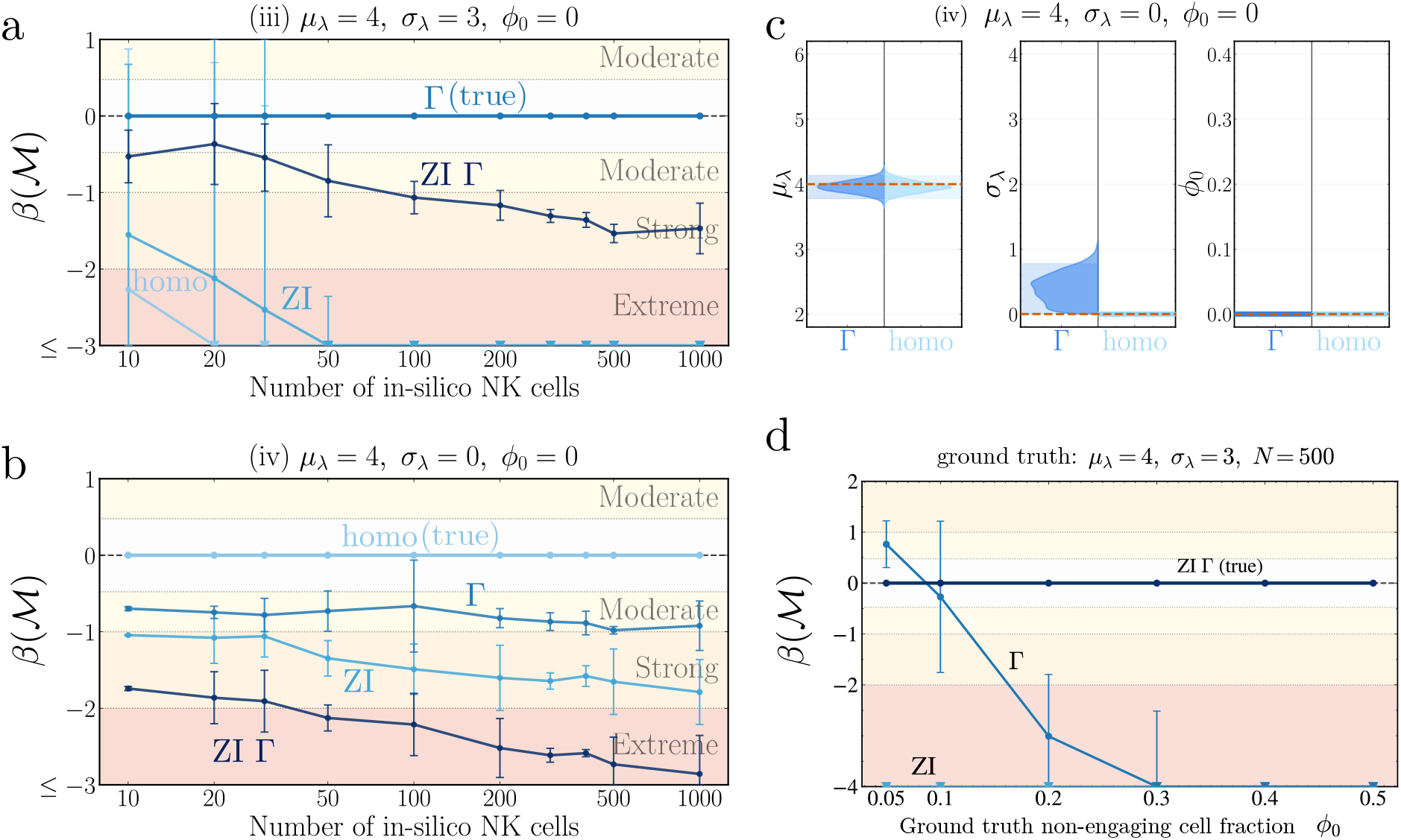
Additional validation of Bayesian inference and model comparison across nested event count population structures. **(a**,**b)** Bayes factor trajectories for synthetic data generated under **(a)** the Gamma model, ℳ_Γ_, with *µ*_*λ*_ = 4, *σ*_*λ*_ = 3, and *φ*_0_ = 0, and **(b)** the homogeneous model, ℳ_homo_, with *µ*_*λ*_ = 4, *σ*_*λ*_ = 0, and *φ*_0_ = 0. All four candidate models were evaluated at each sample size. The vertical axis reports *β*(*ℳ/ ℳ* _true_), such that the true model lies at zero and negative values favour the true model. Points and error bars summarise results across five independent synthetic replicates. Downward triangles indicate values below the displayed range. Background shading denotes conventional strengths of evidence. **(c)** Posterior parameter distributions inferred from synthetic data generated under ℳ_homo_, using the true model, ℳ_homo_, and the second most strongly supported model, _Γ_. Both models recover the ground truth mean event rate *µ*_*λ*_, while the posterior under ℳ_Γ_ concentrates near the homogeneous limit *σ*_*λ*_ = 0. Orange dashed lines denote the ground truth values, and parameters fixed by a model are shown as horizontal lines. **(d)** Sensitivity of model discrimination to the magnitude of zero inflation. Synthetic data were generated under ℳ_ZIΓ_ at *N* = 500, with *µ*_*λ*_ = 4 and *σ*_*λ*_ = 3, while the ground truth zero inflation probability *φ*_0_ was varied.

**Fig. S2.**
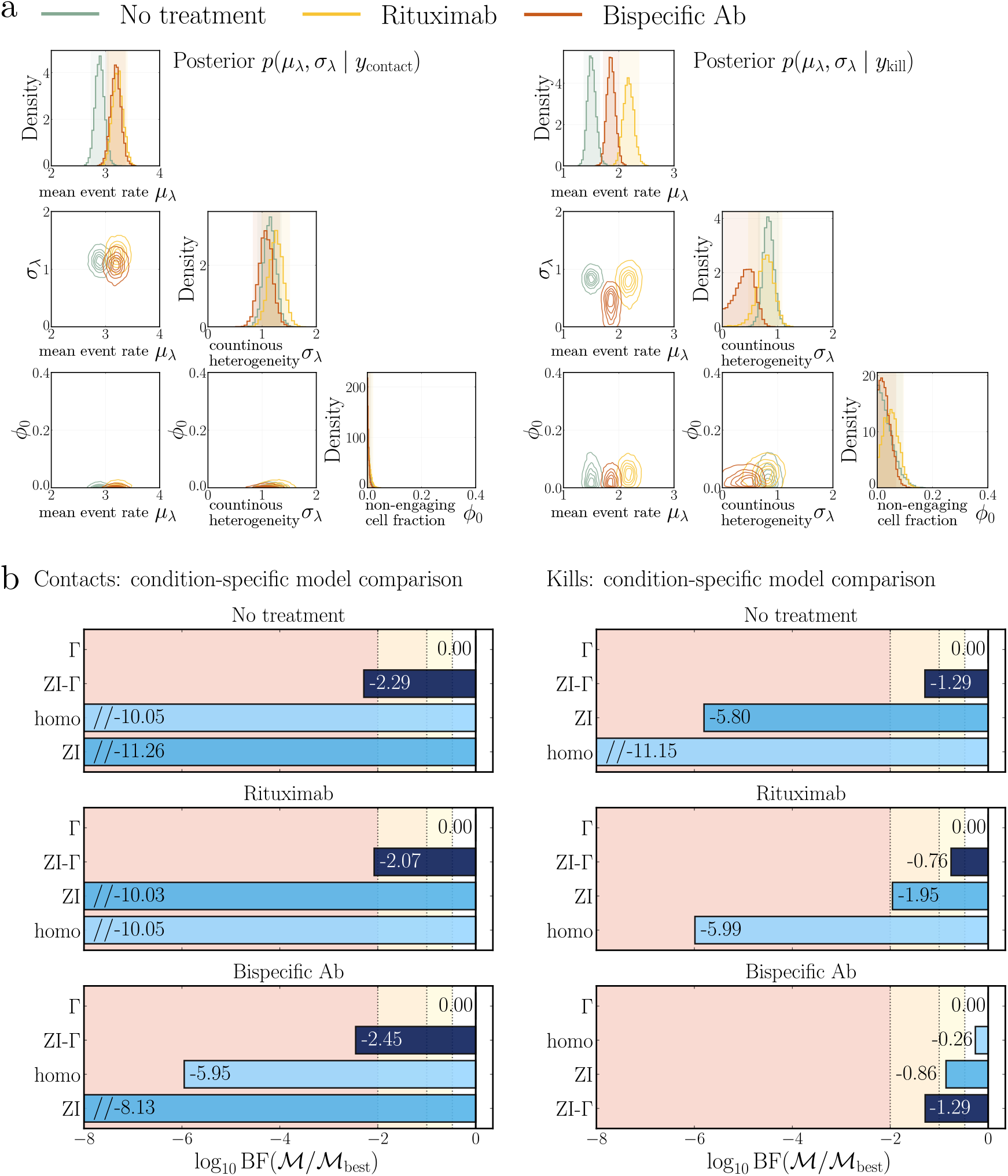
Additional results of Bayesian event-count analysis of NK cell contact and kill distributions. **(a)** Marginal and joint posterior distributions of the mean event rate *µ*_*λ*_ and population heterogeneity *σ*_*λ*_ for contact (left) and kill (right) counts with ℳ_ZIΓ_. **(b)** Condition-specific Bayesian comparison of ℳ_homo_, ℳ_ZI_, ℳ_Γ_, and ℳ_ZIΓ_. Bayes factors are reported relative to the best-supported model.

**Fig. S3.**
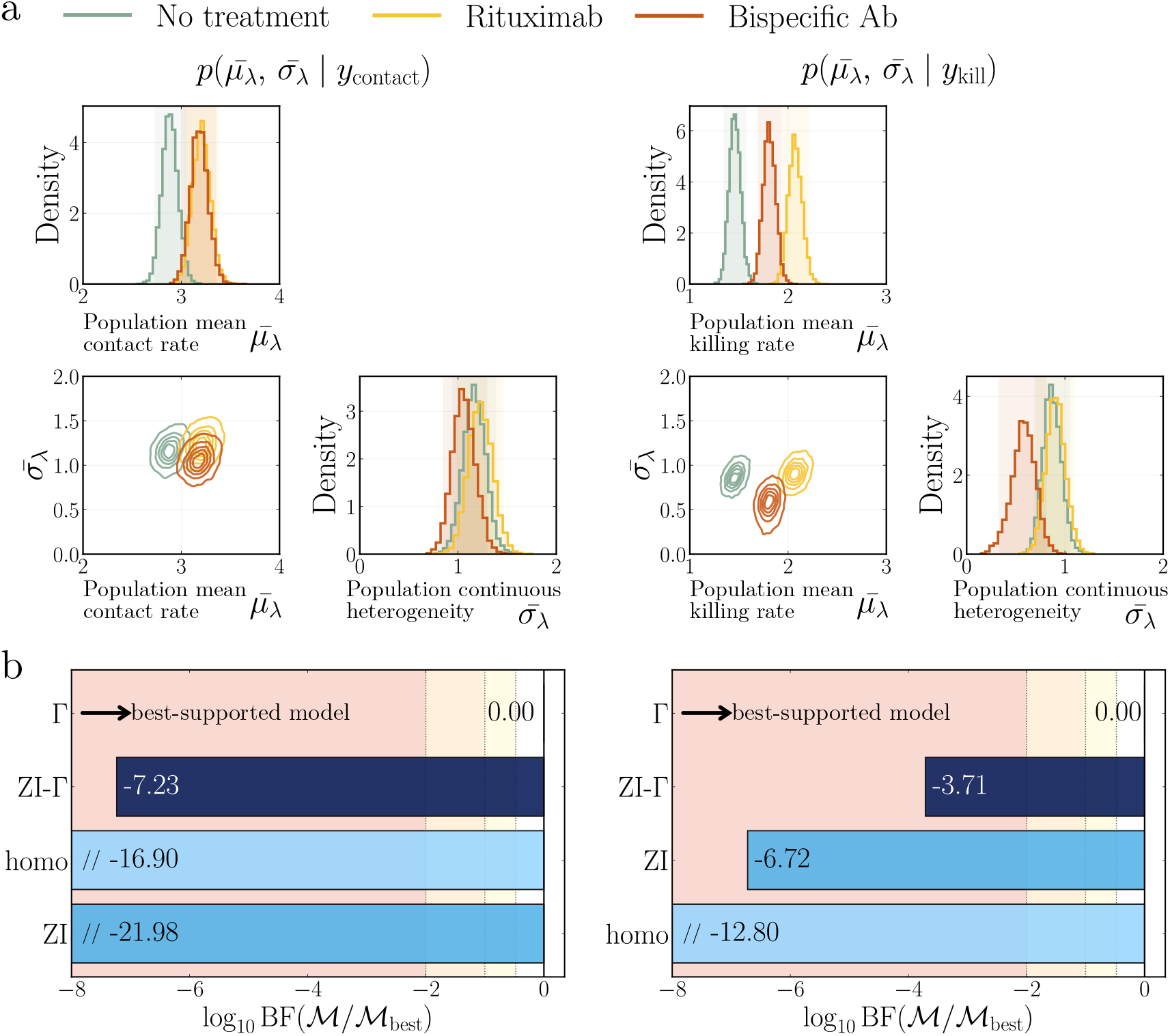
Additional donor-aware Bayesian analysis of NK cell contact and kill counts. **(a)** Marginal and joint posterior distributions of the population mean event rate 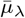 and continuous cell-to-cell heterogeneity 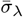, inferred under ℳ_Γ_ for contact counts (left) and kill counts (right). **(b)** Bayesian comparison of the donor-aware ℳ_homo_, ℳ_ZI_, ℳ_Γ_, and ℳ_ZIΓ_ models for contact counts (left) and kill counts (right). Values are log_10_ Bayes factors relative to the best-supported model.

**Fig. S4.**
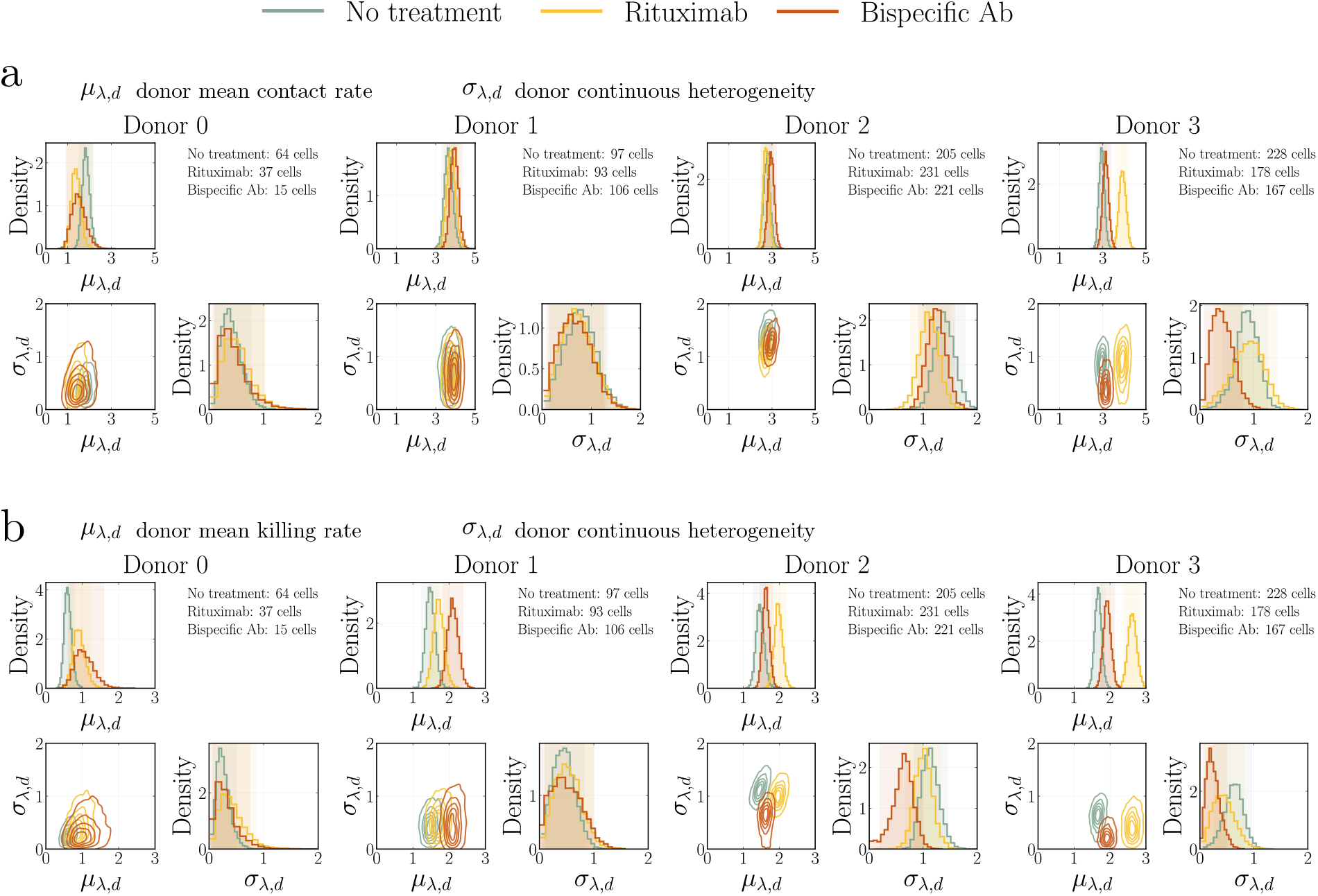
Within-donor comparison of NK cell mean event rates and continuous heterogeneity across treatment conditions. Donor-specific posterior distributions were inferred using the donor-aware ℳ_Γ_ model for **(a)** contact counts and **(b)** kill counts. Results are grouped by donor, with the three treatment conditions overlaid within each donor. Translucent vertical bands indicate the 95% highest density intervals of the corresponding marginal posteriors.

**Fig. S5.**
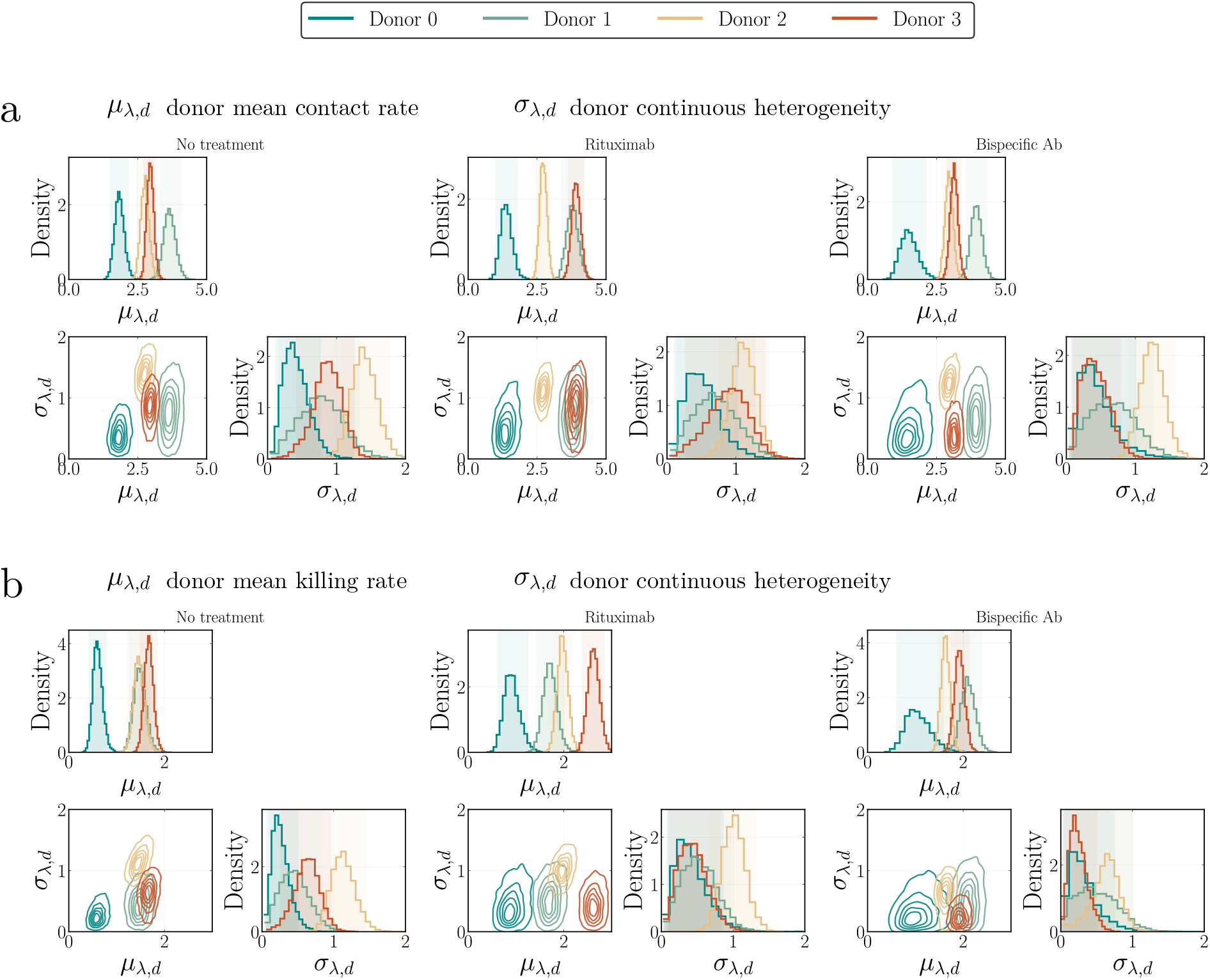
Between-donor comparison of NK cell event rates and continuous heterogeneity within treatment conditions. Donor-specific posterior distributions were inferred using the donor-aware ℳ_Γ_ model for **(a)** contact counts and **(b)** kill counts. Results are grouped by treatment, with the four donors overlaid within each treatment condition. Translucent vertical bands indicate the 95% highest density intervals of the corresponding marginal posteriors.

**Fig. S6.**
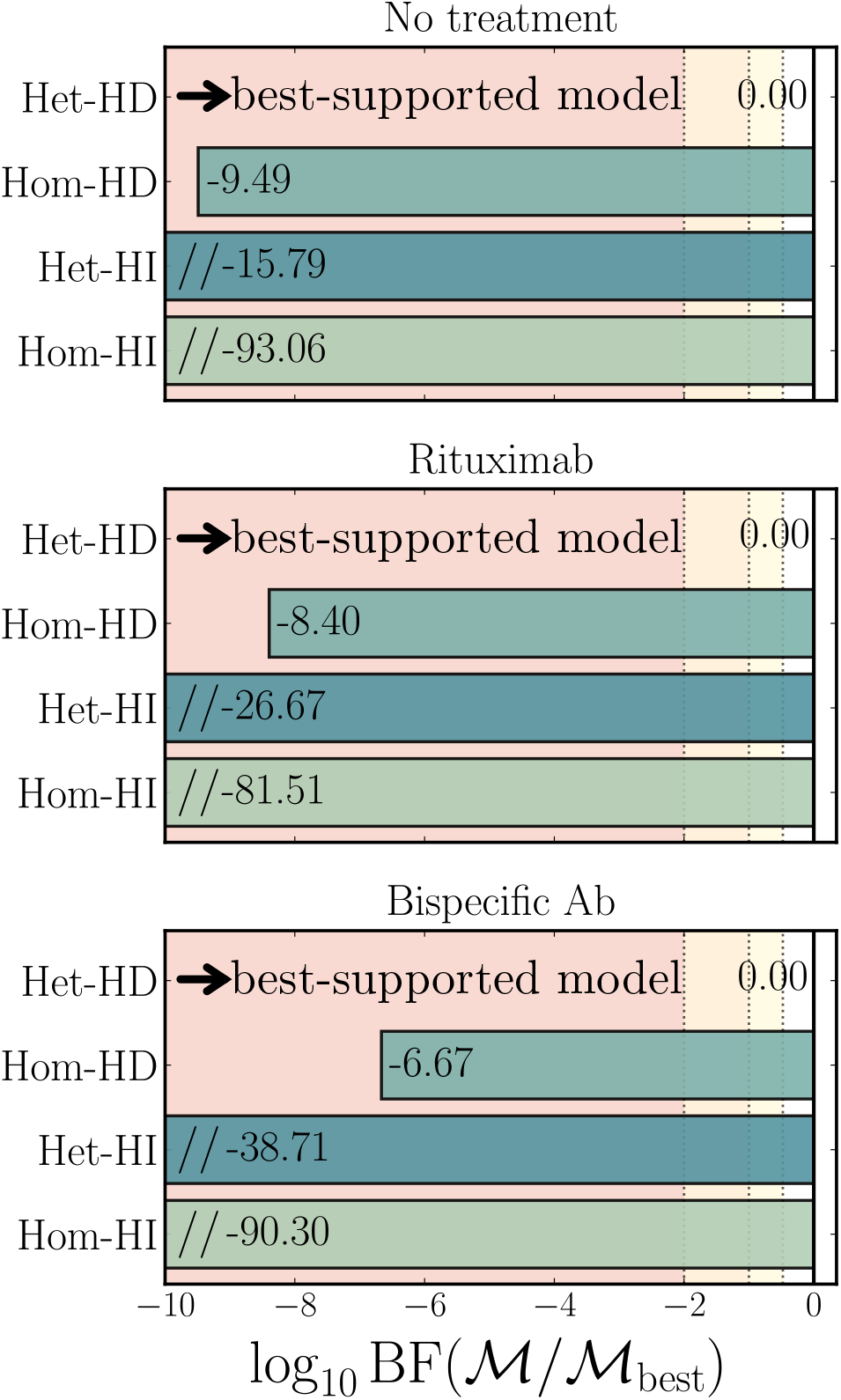
Bayesian comparison of four trajectory models for each treatment condition. Het-HD, heterogeneous history-dependent; Hom-HD, homogeneous history-dependent; Het-HI, heterogeneous history-independent; Hom-HI, homogeneous history-independent. Background shading indicates the strength of evidence against each candidate model: yellow denotes modest evidence, orange denotes strong evidence, and red denotes extreme evidence in favour of the best-supported model.

**Fig. S7.**
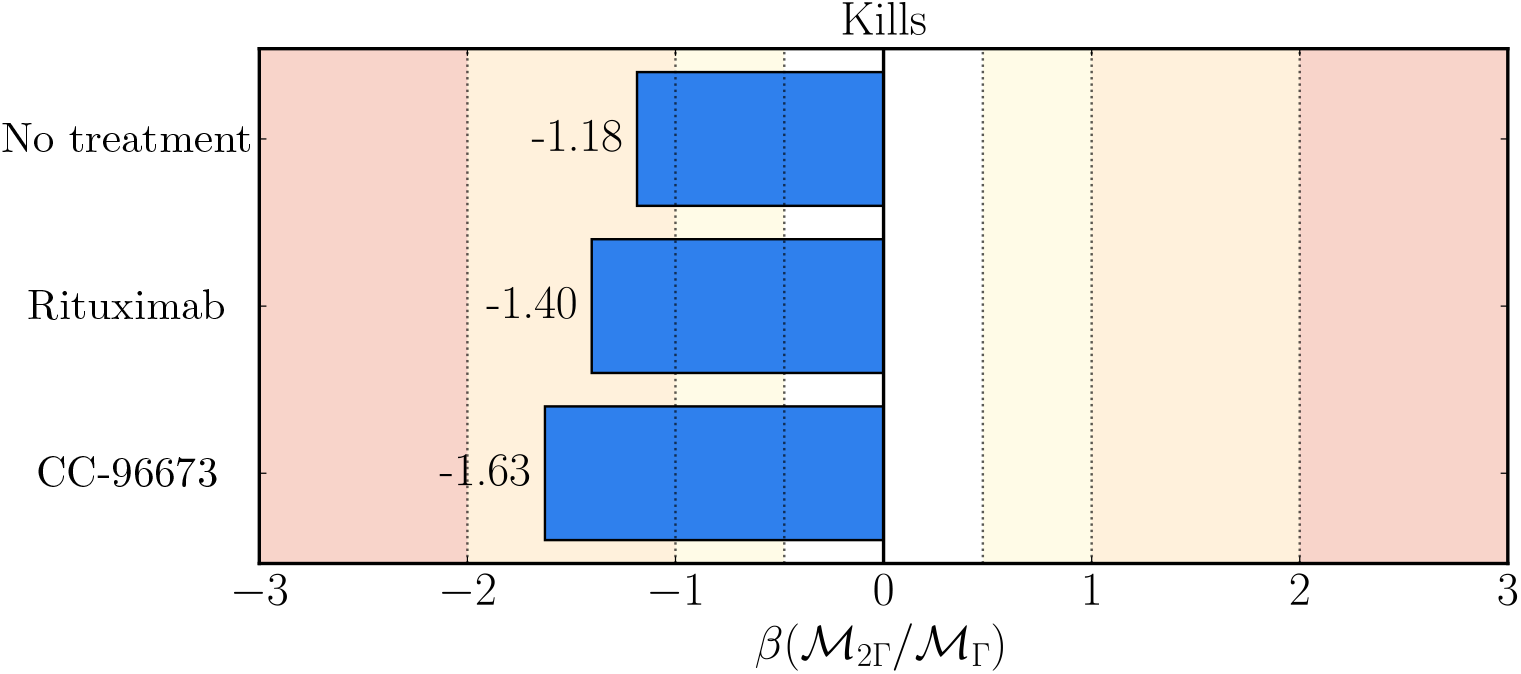
Donor-aware Bayesian comparison of Γ and 2Γ models for kill counts. Bars show *β*(ℳ_2Γ_*/ℳ*_Γ_) = log_10_ BF_2Γ,Γ_, calculated jointly across donors for each condition. Negative values favour ℳ_Γ_, whereas positive values favour _2Γ_. Background shading indicates evidence strength in either direction: yellow denotes modest evidence, orange denotes strong evidence, and red denotes extreme evidence.

**Fig. S8.**
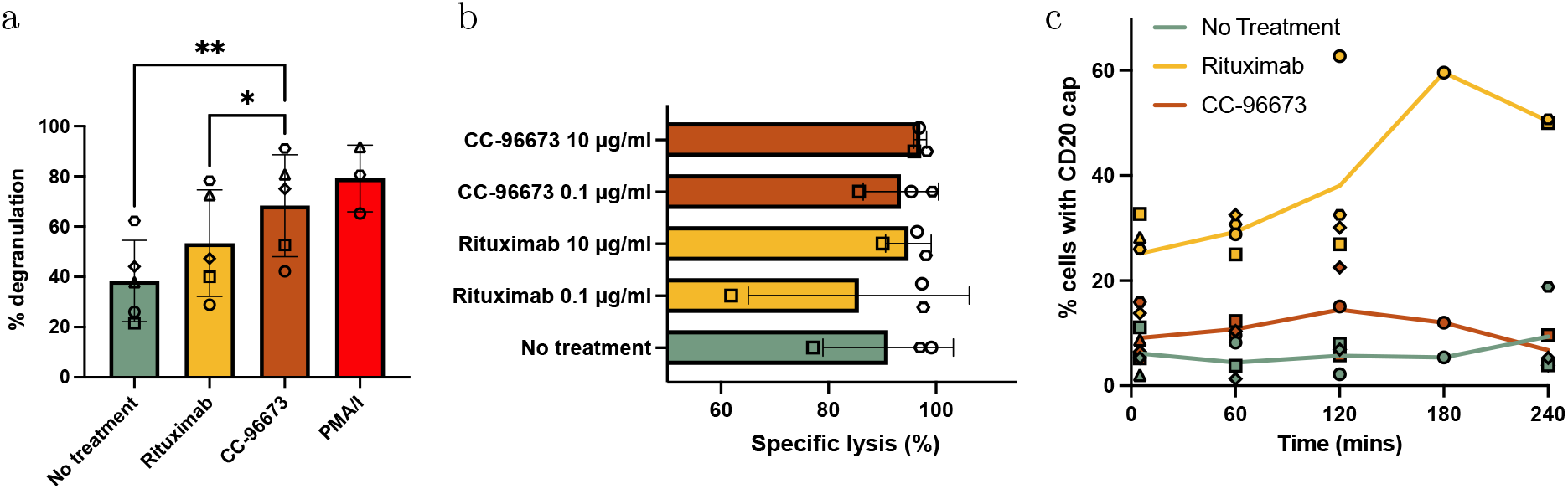
**(a)** NK cell degranulation against the CD20^+^ 721.221 cell line after 4 hours of coculture at an E:T ratio of 1:4. Antibody conditions were tested at 10 µg/mL using five matched donors (*n* = 5). PMA/I was included as a positive control (*n* = 3) and excluded from statistical analysis. A one-way repeated-measures ANOVA with Geisser–Greenhouse correction identified a significant treatment effect (*p* = 0.0032). Dunnett’s post hoc test showed significantly greater degranulation with CC-96673 than with no treatment (*p <* 0.01) and rituximab (*p <* 0.05). Bars show the mean *±* SD, and symbol shapes identify individual donors. **(b)** Specific lysis of 721.221 target cells in the presence of NK cells and the indicated antibody treatments at 0.1 and 10 µg/mL. Cells were incubated for 4 hours at an E:T ratio of 1:4, with three donors per condition (*n* = 3). Bars show the mean *±* SD, and symbol shapes identify individual donors. **(c)** Proportion of 721.221 cells showing a CD20 cap over time following staining with an anti-CD20 mAb (clone L26) and treatment with the indicated antibody at 10 µg/mL. CD20 capping was assessed by confocal microscopy, with at least 50 cells examined per condition in each experimental repeat. Symbol shapes identify individual donors or experimental repeats, and lines connect the mean values at each time point. At least three repeats were performed at the earlier time points. At the final two time points, one repeat was performed at 180 minutes instead of 240 minutes.

**Fig. S9.**
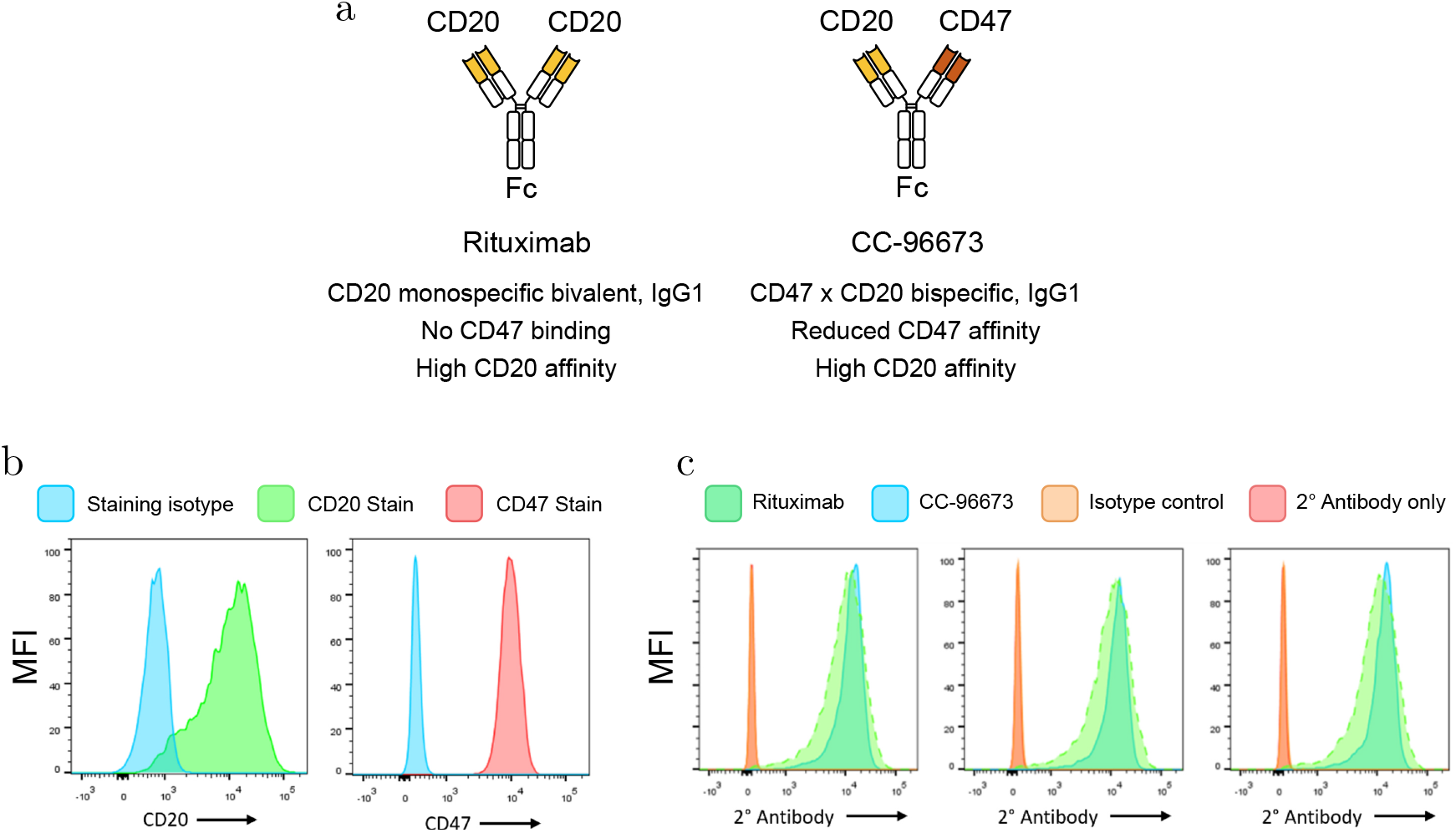
Antibody schematics, 721.221 cell phenotyping and therapeutic antibody binding. **(a)** Comparison of antibodies used. Bispecific on right can bind to CD20 and CD47. Rituximab can bind to CD20 bivalently. Both antibodies are of the IgG1 isotype and have identical Fc regions. **(b)** 721.221 cell lines stained for CD20 and CD47 expression, as well as FMO controls, were acquired on the same day. Marker MFI of live, single cell populations was plotted. **(c)** Mean fluorescence intensity (MFI) of therapeutic antibody binding to 721.221 lines measured by flow cytometry. Therapeutic antibodies were stained with a secondary nanobody. (*n* = 3).

